# Epigenetic regulation of cilia stability and kidney by the chromatin remodeling SWI/SNF complexes

**DOI:** 10.1101/2025.06.10.658863

**Authors:** Xiaoyu Cheng, Qianshu Zhu, Shilin Ma, Xiaoyu Peng, Guanliang Huang, Guifen Liu, Wentao Zhang, Yong Zhang, Cizhong Jiang, Andong Qiu, Ying Cao

**Author notes:** These authors contributed equally (Xiaoyu Cheng, Qianshu Zhu).

## Abstract

Cilia are important subcellular organelles, whose assembly are regulated by master regulator transcription factors including Foxj1 and Rfx proteins. However, whether and how cilia are regulated at epigenetic level remains unknown. We addressed this question by knocking down or knocking out of chromatin remodeling genes. Notably, depletion of multiple components of the switch/sucrose non-fermentable (SWI/SNF) complexes led to ciliopathy-like phenotypes in zebrafish embryos. Specifically, loss of Actl6a, an essential components of the SWI/SNF complexes, resulted in cilia disassembly and cystic kidney, without affectting cilia motility. Omics analyses revealed that in Actl6a-depleted pronephros or embryos, a set of cilia genes—including master regulators *foxj1a* and *rfx2*—were downregulated at transcriptional level, chromatin accessibility level and SWI/SNF binding level. Depletion of *foxj1a* or *rfx2* in zebrafish also caused cilia disassembly and cystic kidney. Furthermore, overexpression of *foxj1a* or *rfx2* mRNA partially rescued the cystic kidney and cilia disassembly phenotypes in *actl6a* mutants. Taken together, our study reveals that the SWI/SNF complexes maintain cilia stability and kidney homeostasis by directly modulating the expression of *foxj1a* or *rfx2*.

## Introduction

Cilia are important organelles involved in cellular motility and environmental signal sensing ^1,2^. They are required for multiple developmental processes, such as left-right asymmetry, kidney development, limb patterning, cardiac development, etc. Defects in cilia structure or function lead to various malformations, including laterality defects, polycystic kidney disease (PKD), obesity, and retinal degeneration, collectively known as ciliopathies^1,3–6^. At the transcriptional level, cilia assembly— especially for motile cilia—is regulated by transcription factors of the Foxj1 and Rfx family members^7^. However, the epigenetic regulation of cilia remains poorly understood.

Gene expression, a fundamental aspect of all biological processes, is tightly regulated. ATP-dependent chromatin remodeling represents a major epigenetic mechanisms that modulates gene expression through nucleosome organization, reposition, ejection or editing, contributing to both gene activation and repression^8–12^. Four subfamilies of ATP-dependent chromatin-remodeling complexes have been identified: imitation switch (ISWI), chromodomain helicase DNA-binding (CHD), switch/sucrose non-fermentable (SWI/SNF), and INO80 complexes^13–17^. Among these, the SWI/SNF complex has well-documented roles in embryonic stem cell (ESC) maintenance, as well as in neurodevelopment, cardiogenesis, myogenesis, and kidney development^18–26^. Nonetheless, it remains unclear whether or how these complexes contribute to ciliary regulation.

Zebrafish serve as an excellent model for investigating ciliary regulation, particularly in the kidney. A genome-wide genetic screen in zebrafish first established a broad association between ciliary dysfunction and PKD^27^. Unlike mammals, which possess only immotile (primary) cilia in the kidney, zebrafish pronephric cilia are motile and consist of both multicilia and single cilia. Single cilia are localized to the distal early tubule, while multicilia are distributed throughout remaining nephron segments in a characteristic “salt and pepper” pattern^28–30^. As ciliary defects lead to PKD in zebrafish, PKD can serve as a reliable readout of normal ciliary structure and function.

To investigate the function of ATP-dependent chromatin-remodeling complexes in cilia biology, we performed a knockdown screen targeting 25 genes encoding chromatin-remodeling subunits. Notably, knockdown of several genes encoding SWI/SNF subunits resulted in cilia-related phenotypes, including body curvature, cystic kidneys, and retinal degeneration. We focused on Actl6a to dissect the role of the SWI/SNF complexes in cilia and kidney development. Phenotypic characterization revealed that Actl6a is essential for maintaining cilia stability and preventing cystic kidney. Furthermore, pronephros-specific RNA sequencing (RNA-seq) analysis and in situ hybridization demonstrated that several cilia-related genes— including the master regulators *foxj1a* and *rfx2*—were downregulated in Actl6a-depleted embryos. Pronephros-specific Assay for Transposase-Accessible Chromatin sequencing (ATAC-seq) revealed reduced accessibility at the regulatory regions of ciliary genes, including *rfx2*, in Actl6a-depleted embryos. In addition, FitCUT&RUN (Fc fragment of immunoglobulin G tagging followed by CUT&RUN) analysis showed decreased SWI/SNF complex binding at the *foxj1a* enhancer in Actl6a-depleted embryos. Overall, cilia genes were significantly enriched among those that were downregulated (RNA-seq), showed reduced chromatin accessibility (by ATAC-seq), and decreased SWI/SNF binding (FitCUT&RUN) in Actl6a-depleted embryos. Knocking out *foxj1a* or *rfx2* also led to cilia disassembly and cystic kidney phenotypes, mimicking the effects of Actl6a loss. Conversely, overexpression of *foxj1a* or *rfx2* mRNA in *actl6a^−/−^* mutants partially rescued both phenotypes. These results suggest that the SWI/SNF complexes maintain ciliary stability and prevent cystic kidney disease through epigenetical regulation of ciliary genes, including *foxj1a* and *rfx2*.

## Results

### The SWI/SNF complex is required to prevent cystic kidney

To investigate the role of chromatin remodeling complexes in cilia regulation and embryonic development, we knocked down 25 genes encoding subunits for the four subfamilies of ATP-dependent chromatin-remodeling complexes individually by morpholinos (MOs), including 10 genes from the SWI/SNF complex (*b*r*g1*/*smarca4a*, *brm*/*smarca2*, *baf250*/*arid1ab*, *smarcc1a*, *arid2*, *baf57*/*smarce1*, *baf60a*/*smarcd1*, *baf53a*/*actl6a*, *snf5*/*smarcb1a*, *brd7*), 3 from ISWI complex (*smarca5*, *baz1a*, *chrac1*), 10 from CHD complex (*chd2*, *chd4a*, *chd4b*, *chd*7, *mta*2, *mta*3, *hdac*1, *parp1*, *rbbp4*, *mbd3a*) and 2 from INO80 complex (*ino80*, *srcap*). Interestingly, knocking down 8 genes individually, including 5 SWI/SNF genes and 3 CHD genes, show cilia-related phenotypes, such as body curvature, cystic kidney and retina degeneration (Supplementary Figure 1A).

By using CRISPR/Cas9 technology, we further generated mutants for three genes with the highest ratio of cystic kidney when knocked down. Mutations in these three genes-*smarca4a*, *smarce1* or *actl6a*-leads to cystic kidney or epicardiac edema (Figure 1B and C, Supplementary Figure 1B). As the *smarca4a* and *smarce1* mutants showed few renal cysts, we speculated that these genes are maternally expressed and the cystic phenotypes are masked by maternal genes, as happened to *pkd2* gene. *Pkd2* is the zebrafish homologue of human *PKD2*, the causal gene for autosomal dominant PKD. Only knockdown, but not knockout, of *pkd2* in zebrafish show cystic kidney due to the maternal deposit of *pkd2* in zebrafish embryos^27^.

**Figure 1:**
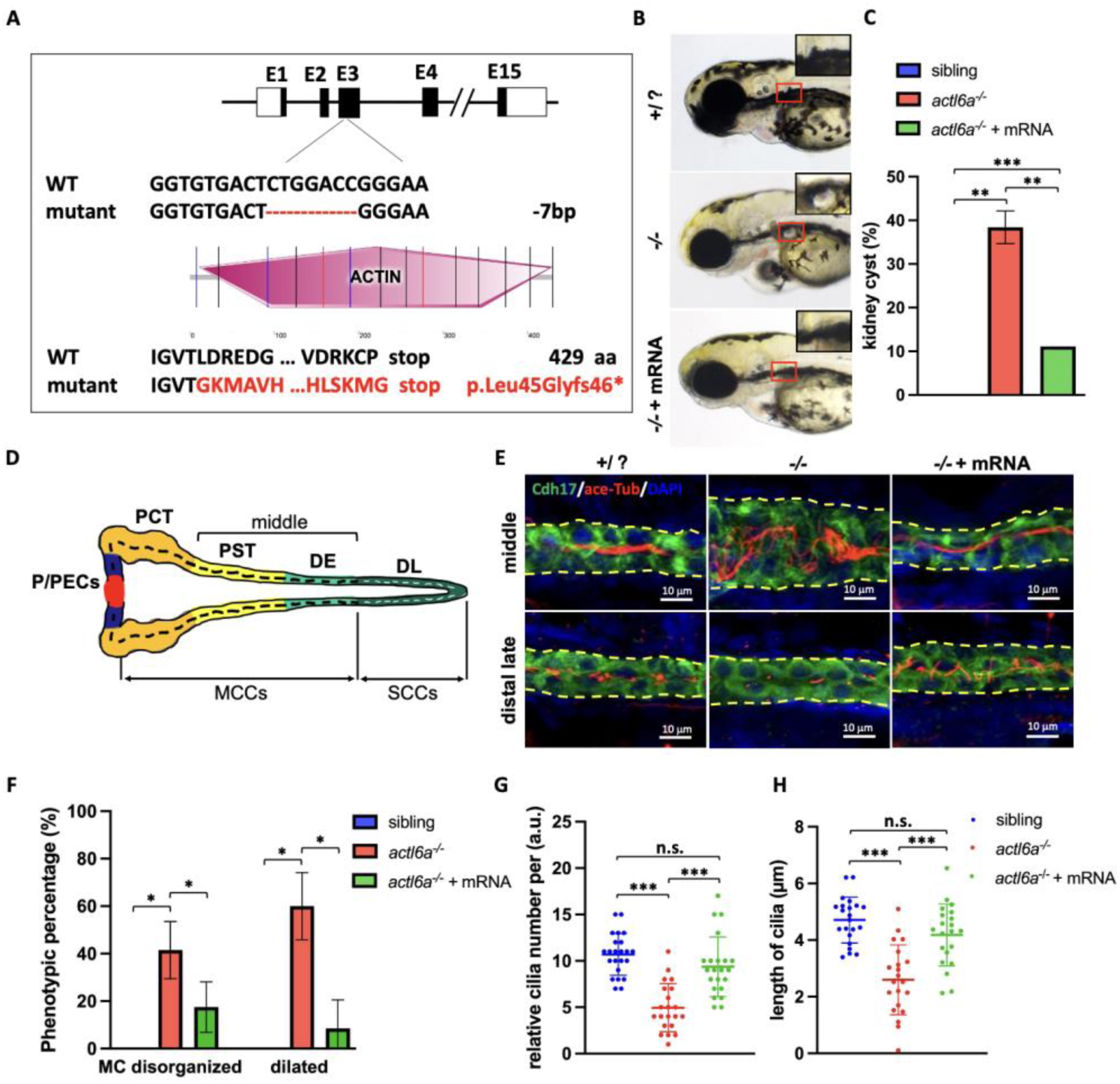
Actl6a is required for ciliary stability and kidney development. (A) Schematic representation of *actl6a* mutant. The gRNA targeting exon 3 induces a 7-base deletion, leading to premature translation termination at 89 aa. Red fonts indicate the sequence with incorrect amino acids. (B) Overexpression of *actl6a* mRNA rescues the phenotype of kidney cysts in *actl6a^−/−^* embryos. Insets show a magnified view of the glomerulus area within the boxed regions. (C) Quantification of the percentage of embryos with kidney cysts in (B). n = 50-150 embryos per group, N = 2 repeats. (D) a schematic of the pronephros in zebrafish embryos at 3 dpf. (E) Immunostaining with antibodies against Acetylated-⍺-Tubulin (red, labels cilia) and Cdh17 (green, labels pnt) shows disorganized multi-cilia in the PST and disassembled single-cilia at the DL of pnt in *actl6a^−/−^*embryos at 3 dpf, which was rescued by overexpression of *actl6a* mRNA. Yellow dotted lines outline the pnt segments. (F) The percentage of phenotypic embryos in the pnt of 3 dpf control siblings, and *actl6a^−/−^*embryos with or without *actl6a* mRNA overexpression, quantified and plotted. n = 8-10 embryos per group, N = 3 repeats. (G-H) Statistics on the number of single cilia per arbitrary unit (a.u.) and cilia length in the distal pnt of 3 dpf control siblings and *actl6a* mutant embryos with or without *actl6a* mRNA overexpression. n = 8-10 embryos per group, N = 2-3 repeats. The data are presented as the mean±SD. (n.s.) not significant, (*) P < 0.05, (**) P < 0.01, (***) P < 0.001. Scale bars, 10 µm. Abbreviations: aa, amino acid; dpf, days post-fertilization; pnt, pronephric tubule; PST, proximal straight tubule; P/PECs, podocytes and parietal epithelial cells; PCT, proximal convoluted tubule; DE, distal early tubule; DL, distal late tubule; MCC, multi-ciliated cell; SCC, single-ciliated cell.

To verify this hypothesis, we conducted in situ hybridization to check the expression patterns of these three genes *smarca4a*, *smarce1* and *actl6a*. Indeed, all three genes are expressed maternally (Supplementary Figure 1C), suggesting that these genes are maternally deposited and that the difference between morphants and mutants of *smarca4a* and *smarce1* may be due to maternal deposition. At 3 days post-fertilization (dpf), *smarca4a*, *smarce1* and *actl6a* were all expressed in the pronephric tubules (Supplementary Figure 1C), supporting their roles in kidney development. Therefore, both functional analysis and expression patterns suggest that the SWI/SNF complex plays a critical role in the kidney development of zebrafish.

### Actl6a is required for cilia stability

To further explore the function of the SWI/SNF complex in kidney development, we focused on Actl6a, one of its essential components. The *actl6a* mutant allele, which carries a 7-bp deletion in the third exon, is predicted to encode a truncated polypeptide with 45 incorrect amino acid residues (p.Leu45Glyfs46*), and is considered a null allele (Figure 1A). The *actl6a* mutants exhibited cystic kidneys with a penetrance of 38.4 ± 3.8%, and overexpression of *actl6a* mRNA significantly reduced the percentage of embryos with cystic kidneys to 11% (Figure 1B and C). Similarly, the cystic phenotype induced by *actl6a* MO could also be rescued by *actl6a* mRNA (Supplementary Figure 1D and E), further confirming that the phenotype was specifically caused by loss of *actl6a*.

To understand the underlying mechanism, we further characterized *actl6a^−/−^*mutant embryos. Immunostaining revealed that, compared with control siblings, the pronephric tubules (pnt) in *actl6a^−/−^* mutant embryos were dilated, and both the number and the length of cilia in the distal pnt were significantly reduced at 3 dpf (Figure 1D-H). These defects could be rescued by overexpression of *actl6a* mRNA (Figure 1D-H), indicating that Actl6a is required for either cilia assembly or stability. To distinguish between these possibilities, we examined cilia morphology at both early (1 dpf, shortly after cilia formation), and later (3 dpf) stages in *actl6a* morphants, since *actl6a* mutants were morphologically indistinguishable from the control siblings. Immunostaining showed that in *actl6a* morphants, cilia number and length were comparable to controls at 1 dpf but markedly reduced at 3 dpf (Supplementary Figure 2B-D), suggesting progressive cilia disassembly rather than a defect in initial assembly. Furthermore, to potential effects of maternally deposited Actl6a, we analyzed cilia in the neuromast cells of the lateral line, where ciliogenesis occus later (around 2 dpf). We found that cilia assembled normally at 2 dpf but were lost by 4 dpf in *actl6a* morphants (Supplementary Figure 2E and F), further supporting a role for Actl6a in cilia maintenance rather than assembly.

We then asked whether other SWI/SNF components are also required for cilia stability. Immunostaining showed that both the number and length of distal cilia were reduced in *smarce1^−/−^* mutants compared with control siblings at both 3 dpf and 4 dpf. Notably, the reduction in cilia number was more pronounced at 4 dpf than at 3 dpf (Supplementary Figure 2G-I), consistent with progressive cilia disassembly. Similarly, in *smarca4a* morphants, cilia number and length were normal at 1 dpf but decreased at 3 dpf (Supplementary Figure 2J-L), indicating cilia disassembly. Together, these findings suggest that the SWI/SNF complex is essential for maintaining cilia stability.

### Cilia genes are down-regulated in the *actl6a-*depleted embryos

To further investigate the mechanism by which Actl6a regulates cilia stability, we performed RNA-seq analysis on the pronephros of *actl6a-*depleted embryos and their wild-type controls at 30 hpf and 72 hpf (Figure 2A). Both heatmap and principal components analysis (PCA) revealed the data sets were highly reproducible, as they were strongly clustered by genotype (Figure 2B, Supplementary Figure 3A-C). A total of 1108 genes were upregulated and 1190 genes were downregulated (|FC|>1.5, p<0.05) in the *actl6a* morphants at 30 hpf, as well as 2007 genes were upregulated and 961 genes were downregulated in the *actl6a* mutants at 72 hpf compared with their control siblings (Figure 2C, Supplementary Figure 3D, Supplementary Table 1). Gene ontology (GO) enrichment analysis using DAVID (https://david.ncifcrf.gov/) revealed that cilia-related terms, such as cilium organization, cilia movement, and cilium assembly, were enriched among the down-regulated genes in *actl6a-*depleted pronephros at both stages (Figure 2D, Supplementary Figure 3E, Supplementary Table 1), suggesting that Actl6a regulates cilia at multiple levels.

**Figure 2:**
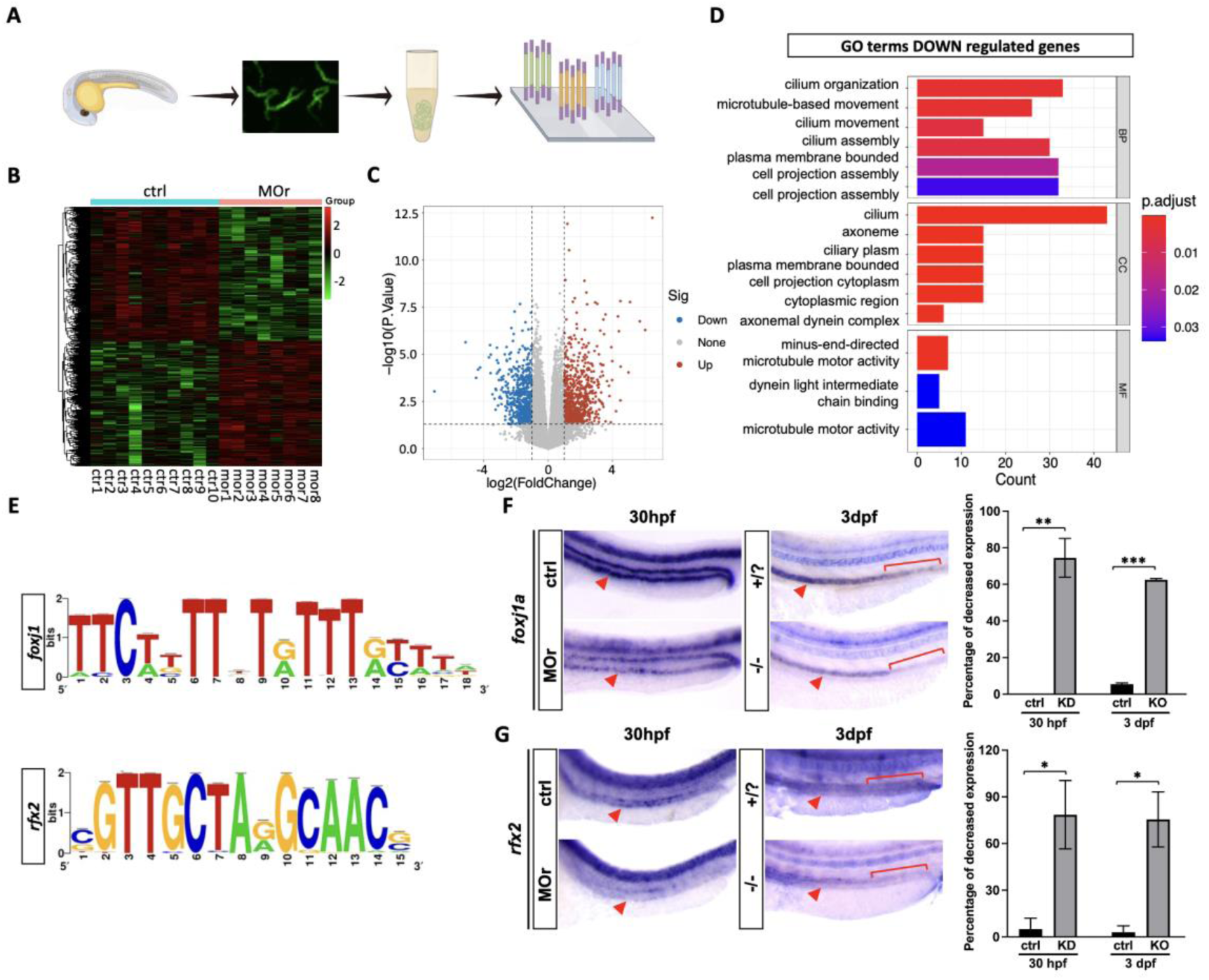
Requirement of *actl6a* for the expression of ciliary genes. (A) Schematic illustration of RNA-Seq library construction for the pronephros of zebrafish embryos. (B) Heatmap of differentially expressed genes of the pronephros between control and *actl6a* morphant embryos at 30 hpf. (C) Volcano plot of RNA-seq results showing differentially expressed genes between *actl6a* morphants and controls at 30 hpf. (D) Gene Ontology (GO) terms of significantly downregulated genes from RNA-seq at 30 hpf. (E) Binding motifs of Foxj1 and Rfx2. (F-G) Whole-mount RNA in situ hybridization (ISH) analysis of *foxj1a* and *rfx2* expression in the pronephric tubule of control, *actl6a* morphant or mutant groups. Red arrowheads indicate pronephric tubules, and the red brackets indicate the distal segments of the pronephros, where expression of the *foxj1a* or *rfx2* is reduced in *actl6a*-depleted embryos. The bar graph on the right represents statistical analysis of ISH results. n = 15-20 embryos per group, N = 2 repeats. The data are presented as the mean±SD. \**P* < 0.05, \*\**P* < 0.01, \*\*\**P* < 0.001. Abbreviations: hpf, hours post-fertilization; dpf, days post-fertilization.

Given that a total of 181 cilia genes were downregulated in the pronephros of *actl6a-*depleted embryos at 30 hpf or 3 dpf, one possibility is that the SWI/SNF complex directly regulates these cilia genes. Another possibility is that the SWI/SNF complex regulates these cilia genes by targeting upstream cilia regulator genes. To explore the latter hypothesis, we used iRegulon to analyze potential regulatory motifs for these cilia genes and found that motifs of several Rfx proteins, as well as Foxj1, were significantly enriched (Figure 2E, Supplementary Table 1). Using ISH, we further examined the expression pattern of both *rfx2* and *foxj1a*, which encode two key ciliary transcription factors. Both genes were found to be downregulated in the pronephric tubule (pnt) of *actl6a*-depleted embryos at 30 hpf or 3 dpf (Figure 2F and G). Notably, both *rfx2* and *foxj1a* were nearly absent from the distal pnt, where single-ciliated cells are located, in *actl6a* mutants at 3 dpf (Figure 2F and G). The reduction of *rfx2* and *foxj1a* expression correlates with cilia disassembly in the pnt of *actl6a*-depleted embryos. These data suggest that the SWI/SNF complex may regulate the expression of cilia genes through targeting *rfx2* and *foxj1a*.

### Cilia motility and structure remain normal in *actl6a-*depleted embryos

Given that the expression of cilia motility genes was downregulated in *actl6a*-depleted embryos (Figure 2D and Supplementary Figure 3E), we hypothesized that cilia motility might be impaired in these embryos. To test this, we examined cilia motility in the pronephric tubules and neural tubes of the zebrafish embryos by using high-speed live imaging. Surprisingly, cilia motility remained normal in both tissues of the *actl6a*-depleted embryos (Figure 3A, Supplementary Figure 4A and B, Supplementary movie 1-6).

**Figure 3:**
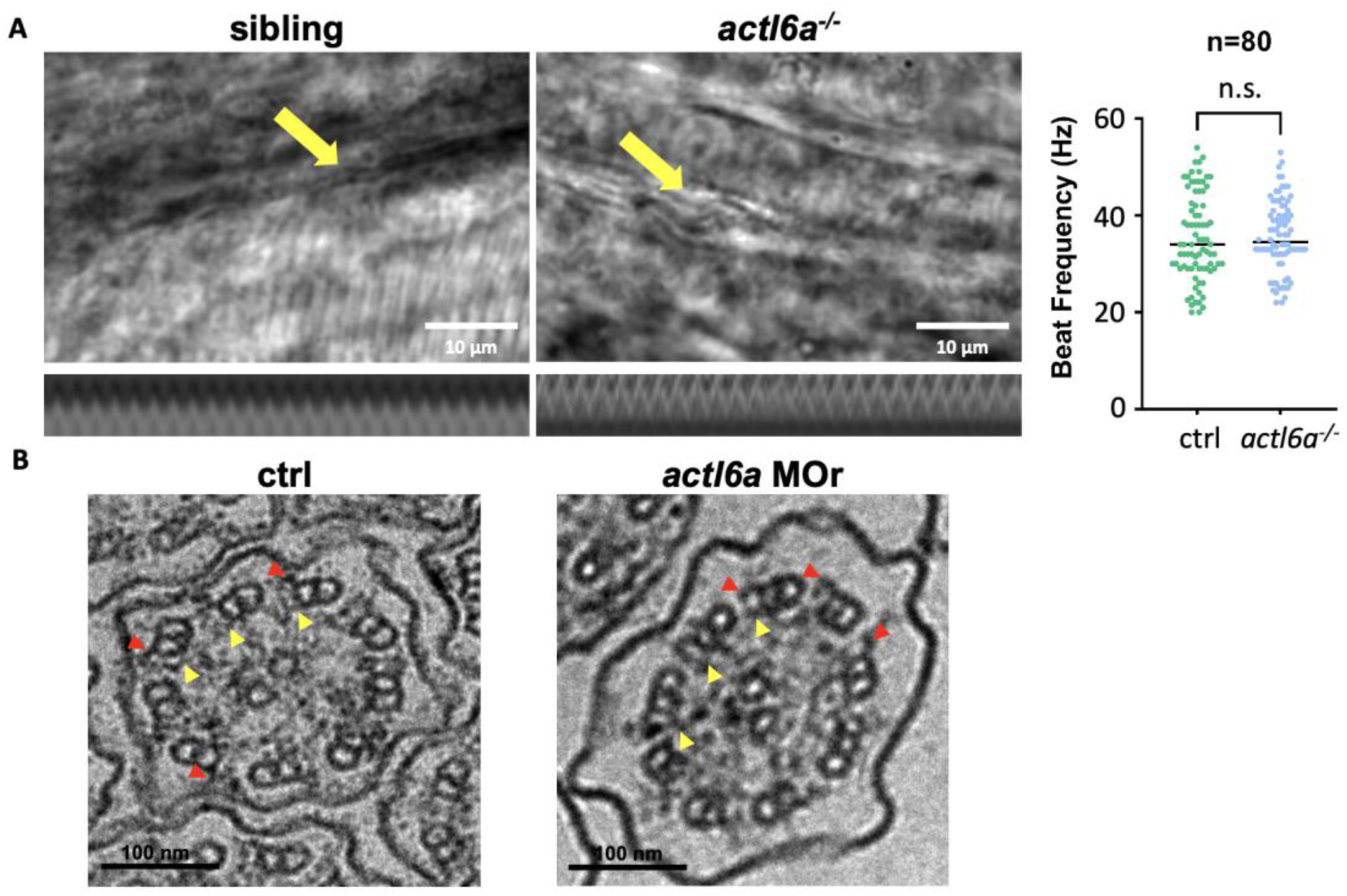
Cilia motility and structure remain normal in *actl6a*-depleted pronephric tubules. (A) Still images showing the beating patterns of multi-cilia in the middle part of pronephric tubules (yellow arrows) in sibling and *actl6a* mutant embryos at 3 dpf. Representative kymographs of ciliary movement are shown below. The right panel shows statistical analysis of ciliary beating frequency. n = 80 (80 cilia from 10 embryos). (B) Transmission electron micrographs show the normal “9+2” microtubule structure of multi-cilia in the middle part of pronephric tubules with clear dynein outer arms (red arrowheads) and dynein inner arms (yellow arrowheads) in both control and *actl6a* morphant groups at 3 dpf. The data are presented as the mean±SD. n.s. not significant. Scale bars, 10 µm in (A), 100 nm in (B). Abbreviations: dpf, days post-fertilization.

We further investigated the ultrastructure of the pronephric cilia by electron microscopy (EM). Consistent with the normal motility, the multi-cilia in the pronephros of *actl6a-*depleted embryos exhibited the same ultrastructure, including the characteristic 9+2 microtubule arrangement, as well as the presence of outer dynein arm (ODA) and inner dynein arm (IDA), as seen in their control siblings (Figure 3B). Thus, both cilia motility and structure remained normal in *actl6a*-depleted embryos, suggesting the SWI/SNF complex regulates cilia stability rather than motility.

### Actl6a regulates chromatin accessibility at the genomic loci of cilia regulator genes

The SWI/SNF complexes are known to control chromatin accessibility to regulate gene expression. To determine the targets of Actl6a, we isolated pronephros from *actl6a* morphants and controls, and performed ATAC-seq analysis (Figure 4A). Principal component analysis (PCA) showed that the experiments were well reproduced (Figure 4B) and the overall chromatin accessibility was significantly affected by Actl6a depletion compared with wild-type controls (Figure 4C). Additionally, the main genomic distribution of differential ATAC-seq peaks was similar in the presence and absence of *actl6a* at promoters (22.53% vs 17.4%) and at potential enhancer sites, such as intronic and intergenic regions (Supplementary Figure 5A). A total of 20,044 accessible peaks were significantly altered, accounting for 67.63% of the total number of accessible peaks across the genome in zebrafish pronephros at 30 hpf. These regulated peaks were associated with 11,444 genes. However, GO enrichment analysis for genes with differential accessibility showed little enrichment for cilia genes (Supplementary Figure 5B). Moreover, among the 7,485 genes with reduced ATAC peaks, only 297 genes were downregulated in our RNA-seq analysis, suggesting the low correlation between accessibility and expression level (Figure 4D), consistent with previous studies^31^. GO enrichment analysis for these 297 genes with both reduced accessibility and down-regulated expression levels revealed enrichment for some ciliary movement genes (Figure 4E). We visualized the ATAC-seq results for selected ciliary genes (*kif3b*, *bbs4*, *dzip1l*, *tmem232*) using the Integrated Genomics Viewer (IGV) software and found that the chromatin accessibility of these cilia genes was significantly reduced (Figure 4H-K). Interestingly, we also observed reduced chromatin accessibility for several genes encoding ciliary transcription factors, including *rfx1a* and *rfx2*. This observation aligns with our ISH results showing that *rfx2* was down-regulated in *actl6a*-depleted pronephros. Additionally, the Actl6a-dependent accessibility peaks corresponded to regulatory regions characterized by H3K27ac and H3K27me3 histone modifications in zebrafish embryos^32^ (Figure 2F and G, Figure 4F and G). These findings suggest that the SWI/SNF complexes may influence the chromatin accessibility of *rfx* genes to regulate the expression levels of cilia genes.

**Figure 4:**
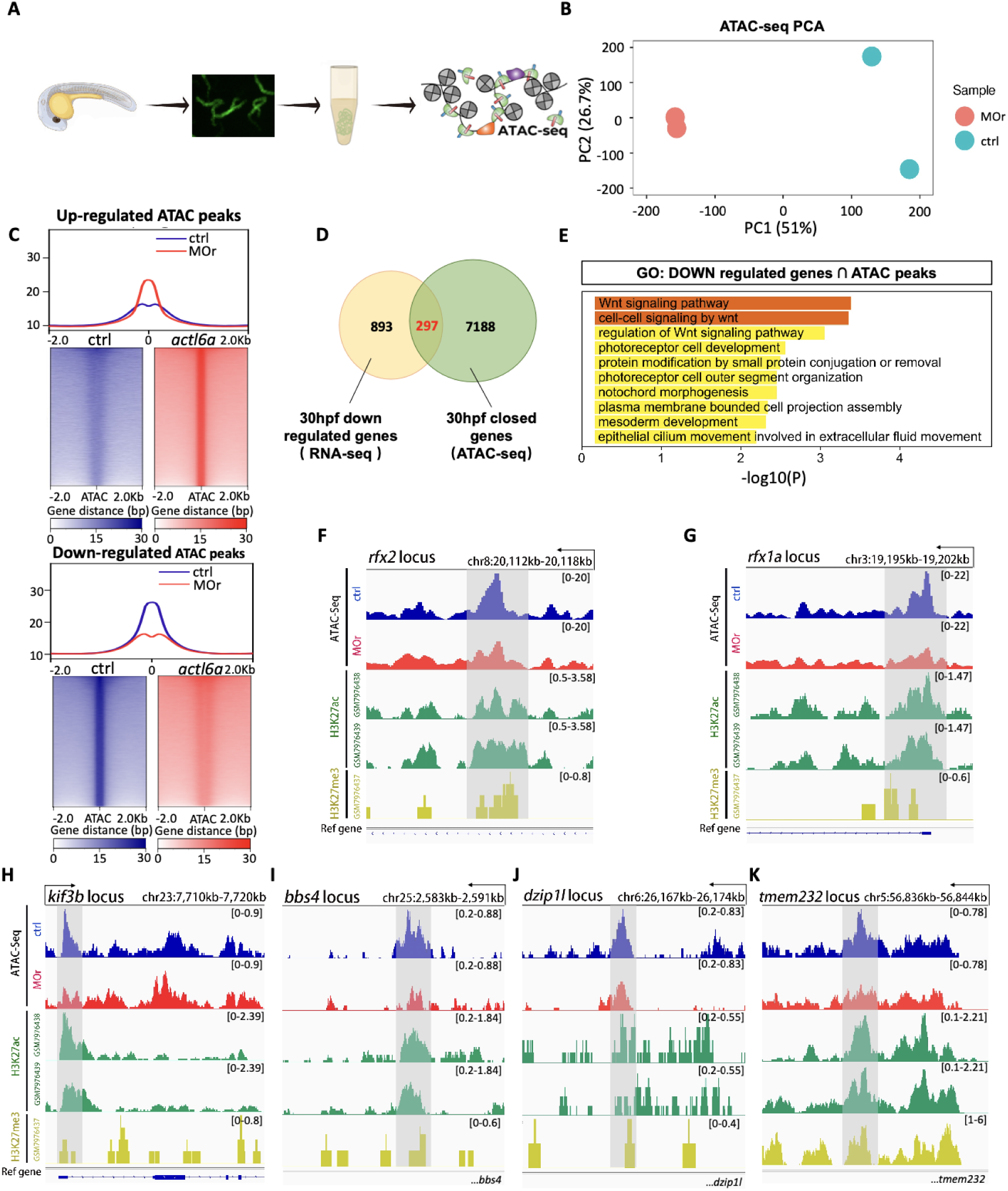
Chromatin accessibility of ciliary genes decreases following *actl6a* depletion. (A) Schematic illustration of ATAC-Seq library construction. ATAC-seq was performed with isolated pronephros from zebrafish embryos at 30 hpf. (B) Principal component analysis (PCA) of ATAC-seq accessibility signals in the pronephric tubules, comparing *actl6a* morphant (red) and control (green) groups. (C) Genome-wide accessibility profiles at upregulated and downregulated ATAC peaks in pronephros of the control and *actl6a* morphants. (D) Venn diagram showing overlap between downregulated genes from RNA-seq (yellow) and closed genes from ATAC-seq (green) in the *actl6a* morphant group compared to the control group at 30 hpf. (E) GO analysis of the intersection gene set of downregulated/closed genes between 30 hpf RNA-seq and ATAC-seq. (F-G) IGV tracks showing decreased accessibility of *rfx2* (F) and *rfx1a* (G) in *actl6a* morphants (red) compared to the control group (blue), overlap with H3K27ac (green) and H3K27me3 (yellow) peaks in *rfx2* enhancer or *rfx1a* TSS from a previously published zebrafish dataset. (H-K) IGV tracks showing reduced accessibility of cilia genes (*kif3b*, *bbs4*, *dzip1l*, *tmem232*) in *actl6a* morphants (red) compared to the control group (blue). The downregulated peaks overlap with enhancer regions of respective genes marked by H3K27ac (green) and H3K27me3 (yellow) peaks from a previously published zebrafish dataset. Abbreviations: hpf, hours post-fertilization; GO, Gene Ontology; IGV, integrative genomics viewer; TSS, transcription start site.

### Cilia genes are targets of the SWI/SNF complexes

Previous studies have shown that Actl6a is not required for the assembly of SWI/SNF complexes, but does regulate the binding of SWI/SNF to genomic regions^33^. To identify the direct targets of Actl6a-regulated SWI/SNF complexes, we applied Fc fragment of immunoglobulin G tagging followed by CUT&RUN (FitCUT&RUN) to *actl6a* knockdown and control embryos (Figure 5A). We first examined the expression of Brg1-Fc, an Fc fragment-tagged version of Brg1, which is a core component of the SWI/SNF complexes, by injecting it into zebrafish embryos. Immunofluorescent staining confirmed that Brg1-Fc was well expressed and localized to the nucleus, as expected, in both *actl6a* knockdown and control embryos (Supplementary Figure 6A).

**Figure 5:**
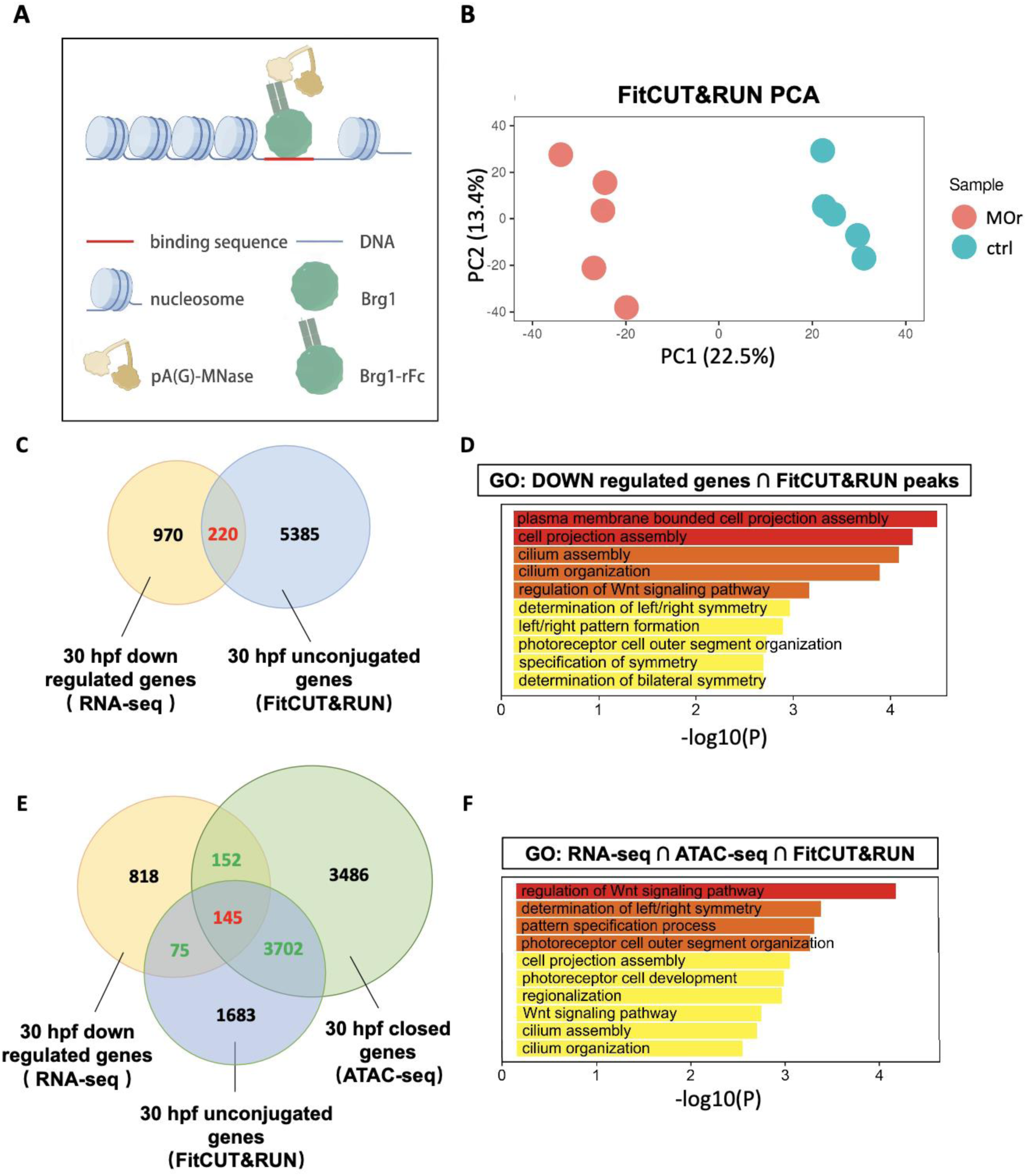
ciliary genes are targets of the SWI/SNF complex. (A) Schematic illustration of the FitCUT&RUN technology. (B) PCA of Brg1 binding signals in zebrafish embryos comparing *actl6a* morphant (red) and control (blue) groups. (C-D) Venn diagram (C) and GO analysis (D) of the intersection gene set between downregulated/unconjugated genes from RNA-seq (yellow) and FitCUT&RUN (blue) in the *actl6a* morphant group compared to the control group at 30 hpf. Note cilia-related GO terms are enriched in (D). (E) Venn diagram showing overlap between downregulated/closed/unconjugated genes from RNA-seq (yellow), FitCUT&RUN (blue), and ATAC-seq (green) in the *actl6a* morphant group compared to the control group at 30 hpf. (F) GO analysis of the intersection gene list (145 genes) from (E). Abbreviations: PCA, Principal component analysis; GO, Gene Ontology; hpf, hours post-fertilization.

PCA analysis showed that the experiments were well reproduced (Figure 5B). We identified 5,605 genes that were unbound and 6,449 genes that were bound by SWI/SNF complexes in the *actl6a* morphants compared with the control embryos (Supplementary Figure 6B and C). GO analysis of these genes with altered SWI/SNF binding showed limited enrichment for cilia-related terms (Supplementary Figure 6D). However, the binding of *foxj1a* by SWI/SNF complexes was reduced, and this unbound region was enriched with H3K27ac or H3K27me3^32^ (Supplementary Figure 6E), suggesting that the altered binding region was a regulatory enhancer region. Moreover, for 220 Actl6a-dependent SWI/SNF target genes, which were both down-regulated (RNA-seq) and unbound by SWI/SNF complexes (FitCUT&RUN), GO analysis revealed enrichment for cilia-related terms, including cell projection assembly, cilium assembly, and left/right pattern formation (Figure 5C and D). This suggests that cilia genes are direct targets of Actl6a-associated SWI/SNF complexes. Interestingly, 145 of these 220 SWI/SNF target genes also showed reduced chromatin accessibility in *actl6a* morphants, with GO analysis showing enrichment for cilia-related terms (Figure 5E and F, Supplementary Table 2). This supports the role of SWI/SNF complexes in regulating the expression of cilia genes by modulating chromatin accessibility. More intriguingly, among these 145 genes that were down-regulated (RNA-seq), had reduced-accessibility (ATAC-seq), and unbound (FitCUT&RUN) by SWI/SNF complexes in *actl6a*-depleted embryos, 21 are known binding targets of Foxj1a or Rfx2^34,35^ (Supplementary Figure 7). This suggests that Actl6a-associated SWI/SNF complexes may work in conjunction with Foxj1a or Rfx2 to regulate the expression of cilia genes.

### Both Foxj1a and Rfx2 act downstream of SWI/SNF complexes to regulate cilia stability

Previous studies have shown that Foxj1a and Rfx2 regulate cilia assembly, particularly for motile cilia^7,36,37^, but their roles in cilia stability have not been addressed. Using CRISPR/Cas9 technology, we knocked out *foxj1a* or *rfx2.* Both crispants (embryos injected with gRNA and Cas9 ribonucleoprotein complex) exhibited heart edema and kidney cysts (Figure 6A and B). Immunostaining revealed that pronephric tubules were dilated, multi-cilia were disorganized, and the number of single cilia were significantly reduced in both *foxj1a* and *rfx2* crispants at 3 dpf (Figure 6A and B, Supplementary Figure 8A and B). However, the single cilia in the pnt were relatively normal in both *foxj1a* and *rfx2* crispants at early stage (1 dpf) (Supplementary Figure 8C and D), suggesting that both Foxj1a and Rfx2 play roles in regulating cilia stability. These findings indicate that both *foxj1a* and *rfx2* crispants phenocopied *actl6a-*depleted embryos in terms of cilia disassembly. Furthermore, overexpression of *foxj1a* and *rfx2* mRNA rescued the cystic kidney and cilia stability defects in a significant proportion of *actl6a^−/−^* mutant embryos (Figure 6C-F), supporting the idea that Actl6a-associated SWI/SNF complexes regulate cilia disassembly and pronephros development through Foxj1a and Rfx2.

**Figure 6:**
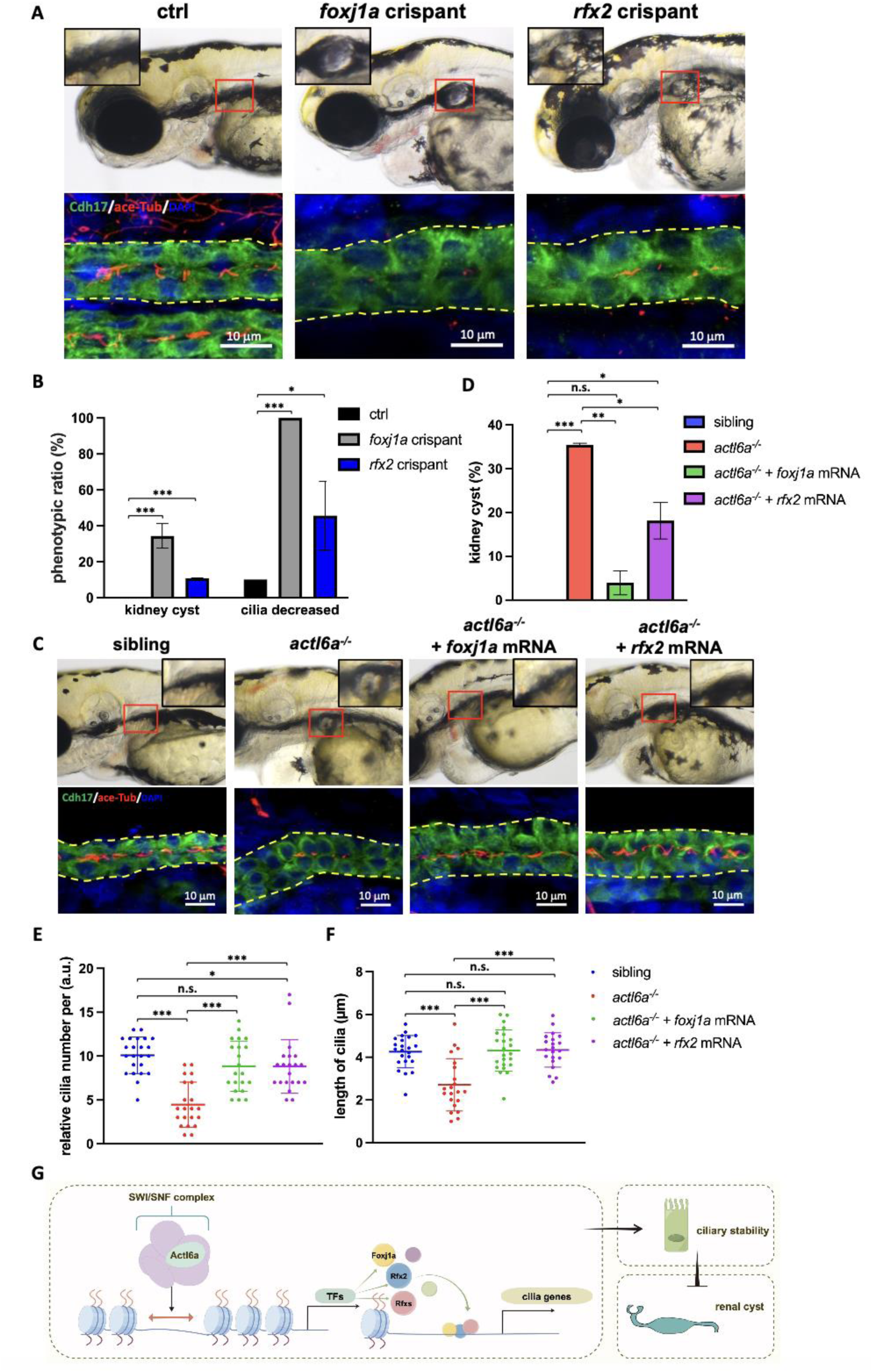
Actl6a regulate ciliary stability through Foxj1a and Rfx2 in the zebrafish pronephros. All embryos are at 3 dpf. (A) The upper row of images shows the lateral views of control embryos, *foxj1a* and *rfx2* crispants. Insets show a magnified view of the glomerulus area within the boxed regions. Note the kidney cysts in *foxj1a* and *rfx2* crispants, but not in control embryos. The bottom row of images shows disassembled single-cilia (Acetylated-⍺-Tubulin, red) in the distal late of the pronephros (Cdh17, green) in the *foxj1a* and *rfx2* crispants but not in the control embryos. Yellow dotted lines outline the pnt. n = 8-10 embryos per group, N = 2 biological repeats. (B) The percentage of renal cyst and decreased single-cilia in the pronephric tubules of crispants is quantified and plotted. n = 50-150 embryos per group, N = 2-3 biological repeats. (C) The representative images show the lateral views of live embryos (upper row) or cilia (Acetylated-⍺-Tubulin, red) in the distal pronephric tubules (bottom row) in *actl6a* mutant embryos with or without overexpression of indicated mRNA and control siblings. Overexpression of *foxj1a* or *rfx2* mRNA rescue both cystic kidney and cilia disassembly defects in *actl6a* mutant embryos. Insets show a magnified view of the glomerulus area within the boxed regions, and yellow dotted lines outline the pnt. Data are quantified and plotted in (D-F). (D) n = 50-150 embryos per group, N = 2 repeats. (E-F) n = 8-12 embryos per group, N = 2 repeats. (G) Schematic illustrating the function of the SWI/SNF chromatin remodeling complex in maintaining ciliary stability and preventing cystic kidney. SWI/SNF complexes bind and modulate the accessibility of key transcriptional regulators of the ciliogenesis program, including *foxj1a*, *rfx2*, etc., facilitating the expression of these transcription factors. Additionally, SWI/SNF complexes regulate chromatin accessibility at genomic loci of ciliary gene, enhancing transcription factor binding and promoting the expression of cilia-related genes. This figure was created with Figdraw (https://www.figdraw.com/<u>#/</u>). The data are presented as the mean±SD. n.s. not significant, \**P* < 0.05, \*\**P* < 0.01, \*\*\**P* < 0.001. Scale bars, 10 µm. Abbreviations: dpf, days post-fertilization; pnt, pronephric tubule.

## Discussion

In this study, we elucidated the role of the SWI/SNF complexes in regulating cilia stability and kidney development. We discovered that 5 out of 10 genes encoding SWI/SNF subunits are essential for normal pronephros development, as knockdown of these genes led to cystic kidney phenotypes in zebrafish. Further characterization of *actl6a*-depleted embryos showed that SWI/SNF complexes are specifically required for cilia stability and pronephros development. Multi-omics analysis revealed that a set of cilia genes, including *foxj1a* and *rfx2*, were downregulated at the transcriptional, chromatin accessibility and SWI/SNF binding levels, indicating that SWI/SNF complexes regulate these cilia genes either directly or by modulating the expression of *foxj1a* and *rfx2*.

There is substantial evidence supporting that SWI/SNF complexes directly regulate *foxj1a* and *rfx2*. In *actl6a*-depleted embryos, both *foxj1a* and *rfx2*, along with their target genes, were downregulated. Chromatin accessibility of the promoter regions for several *rfx* genes, including *rfx2*, was reduced, and the binding of SWI/SNF complexes to the genomic region of *foxj1a* was also decreased. Consistent with our findings, a recent study reported that both GEMC1 and MCIDAS, which are crucial for the expression of *foxj1* and *rfx2*, interact with SWI/SNF components and that these complexes are necessary for the induction of *rfx2* by GEMC1 and MCIDAS^38^. Together, our functional and multi-omics data suggest that SWI/SNF complexes directly regulate the expression of *foxj1* and *rfx2*, thereby influencing cilia stability (Figure 6G).

Cilia are critical for normal kidney development and function in vertebrates. While the cilia in mammalian kidneys are primary/immotile cilia, both multi-cilia and single cilia in zebrafish pronephros are motile, as evidenced by ultrastructure and live imaging studies, consistent with previous reports^39^. Ultrastructural analysis shows that pronephric cilia maintain a “9+2” structure with both ODA and IDA (Supplementary Figure 4C). Live imagining confirms that both multi-cilia and single cilia are motile (personal communication with Prof. Chengtian Zhao). Loss of *actl6a* leads to the disassembly of single cilia in the distal pronephric tubule and results in cystic kidneys, suggesting the essential role of single cilia in kidney development. Notably, previous studies have also shown that loss of single-cilia leads to cystic kidneys in both embryonic and adult zebrafish^40^. Here we provide additional evidence that distal single cilia are crucial for preventing cystic kidney formation in zebrafish embryos. We propose that the single cilia in zebrafish pronephros are functionally equivalent to primary cilia in mammal kidneys, as both play critical roles in preventing cystic kidney formation.

Furthermore, Rfx2 and Foxj1 are key regulatory factors in ciliogenesis, controlling the expression of various genes involved in cilia assembly and structure, including members of the IFT family and BBS families^7,35,41^. However, to our knowledge, their roles in cilia stability have not been explored. IFT particles transport structural proteins, such as tubulin, to the distal end of the flagellum for integration, compensating for subunits removed during disassembly in the turnover process^42–44^. In our study, IFT and BBS family genes, including *ift88*, *ift140*, *ift172*, *bbs2*, *bbs4* and *bbs7*, were significantly downregulated in the *actl6a*-depleted embryos. This suggests that Actl6a controls the expression of downstream ciliary genes by regulating *rfx2* and *foxj1*, thereby ultimately maintaining cilia stability.

By varying its subunit composition, the SWI/SNF chromatin remodeling complex can form hundreds of distinct assemblies, each regulating specific biological processes ^45–47^. Our genetic analyses have revealed that several key subunits, including Smarca4a/Brg1, Smarce1, Smarca2, Actl6a, and Smarcb1a, are essential for kidney development, likely by modulating cilia stability. These findings highlight the critical role of SWI/SNF in maintaining both ciliary function and renal architecture. Although our study provides a foundational understanding of how SWI/SNF complexes contribute to kidney development, further investigation is required to fully elucidate their broader functions in cilia and nephrogenesis.

In conclusion, our work presents a mechanistic model in which SWI/SNF complexes regulate cilia stability and kidney development by controlling chromatin accessibility and the expression of key ciliary genes. These insights not only deepen our understanding of SWI/SNF-related renal phenotypes observed in disorders like Coffin-Siris syndrome but also open up potential therapeutic avenues for ciliopathies and renal malformations associated with chromatin remodeler mutations.

## Materials and Methods

### Zebrafish husbandry

Wild Tubingen zebrafish were bred according to the standard protocol^48^. The previously reported transgenic line *Tg (Na,K-ATPase alpha1A4:GFP)* in which GFP is highly expressed in pronephric epithelia was obtained from Iain Drummond Lab^28^. The zebrafish embryos for injection were obtained through natural mating and cultured at 28.5°C.

### Microinjection

All MOs, gRNAs or mRNAs were injected into zebrafish embryos at the 1-cell stage. Antisense MO oligos were all synthesized by GeneTools (USA).

MOs were used to block the translation of the following genes: *smarca4a* (5′-CATGGGTGGGTCAGGAGTGGACATC-3′), *smarce1* (5′-GGGCCGCTTTGACATCTTGATTGTA-3′), *smarca2* (5′-GGCCATCTATCAGATGAGAATCTTC-3′), *actl6a* (5′-CGTAGACTCCGCCACTCATTTTTAC-3′), *smarcb1a* (5′-TGTCTTGCTTAAAGCCATGTTTGAC-3′), *ift88* (5′-CTGGGACAAGATGCACATTCTCCAT-3′), and *ift20* (5′-CAACAACGTACCTTCATTTTTTCAG-3′). A standard control MO (5′-CCTCTTACCTCAGTTACAATTTATA-3′) was used as a negative control. The concentrations of MOs used for injection were as follows: 0.15 mM for *smarca4a* MO, 0.75 mM for *smarce1* MO, 3 mM for *smarca2* MO, 1.5 mM for *smarcb1a* MO, 1 mM (full concentration) and 0.25 mM (hypomorphic concentration) for *actl6a* MO, 0.05 mM for *ift88* MO, and 0.5 mM for *ift20* MO.

For mRNA injection, C-terminal Fc-tagged *smarca4a*, GFP-tagged *actl6a*, *foxj1a*, *rfx2*, *smarce1* and *smarca4a* constructs were cloned by PCR and then inserted into pXT7 vectors. The mRNA was in vitro transcribed with the mMESSAGE mMACHINE™ T7 Kit (Ambion, USA) using the linearized construct as a template.

### Mutagenesis by CRISPR/Cas9 system

The CRISPR/Cas9 system was applied to generate the zebrafish mutants as previously described^49^. In brief, 1 nL of the mixture of guide RNAs (gRNAs) (100 ng/µL) and Cas9 protein (4 µm) was injected into 1-cell stage embryos. After injection, the target sequences (covering the gRNA target) were PCR amplified from genomic extracted from the injected embryos at 1 dpf. The PCR products were then digested by restriction enzymes to estimate the mutagenesis efficiency. The injected embryos were raised to adulthood and outcrossed to wild-type fish. Positive founder fish (F0) were identified by genotyping the offspring embryos from these outcrosses. The sequences of the gRNA and primers for genotype identification are provided in Reagents table. F1 fish, offspring of the positive F0 fish, were raised and genotyped to obtain stable mutant lines. The live embryos were photographed with a stereo microscope (SMZ1500; Nikon, Japan).

### *In situ* hybridization

Whole-mount in situ hybridization (ISH) followed previously described procedures^50^. Antisense probes for *actl6a*, *smarce1*, *smarca4a*, *foxj1a* and *rfx2* were synthesized using dUTP (Roche, Switzerland), T7 RNA polymerase (NEB, USA) and PCR-cloned DNA templates. The stained embryos were dehydrated in glycerol and photographed with a stereo microscope (SMZ1500; Nikon, Japan).

### Immunohistochemistry

For histological observation of cilia in the pronephros, the embryos were fixed in Dent’s fixative overnight at −20°C. Primary antibodies were diluted in PBST (PBS with Tween 20) containing 10% donkey or goat serum at the following concentrations: goat anti-GFP (1:500; Rockland, USA) and mouse anti-acetylated tubulin (1:2000; Sigma, USA). Rabbit anti-Cdh17 (1:300) was generated by B&M. The embryos were photographed using a confocal laser microscope (SP8; Leica, Germany).

For BRG1-lFc staining, the 30-hpf embryos were fixed in 4% formaldehyde overnight at 4°C and then permeabilized in ice-cold acetone for 7 min at −20°C. After direct staining with 488 goat anti-rabbit IgG (H&L) (1:500; Thermo Fisher Scientific, USA), they were photographed using a confocal laser microscope (SP8; Leica, Germany).

Cilia number and length were measured using Image J (National Institutes of Health, USA) for 6-10 different embryos per group. The confocal images showing discernable individual cilia base and ends were outlined, and the calculated length was recorded.

### Transmission electron microscopy

The embryos were prepared for electron microscopy according to previously published protocols^51^. Briefly, embryos were fixed with 2% paraformaldehyde and 2.5% glutaraldehyde in 0.1 M phosphate buffered saline (PBS) for 2 hours at 37°C. Embryos were then washed three times with 0.1 M PBS and treated with tannic acid at room temperature for one minute. The embryos were placed in 2% osmium tetroxide and left at 4°C for 1 hour, and then washed with PBS. Subsequently, the liquid was replaced with 1% uranyl acetate and incubated at 4°C for 1 hour. Embryos were dehydrated through a series of 20%, 30%, 40%, 60%, 80% and 100% ethanol, and finally in acetone to infiltrate with embed 812 resin. Embryos were embedded in 100% 812 resin and polymerized at 60°C overnight. Finally, ultra-thin sections were cut on an EM UC6 ultramicrotome (Leica, Germany) and examined with a transmission electron microscope (JEM-1230; JEOL, Japan).

### High-speed video microscopy

Ciliary motility was recorded in the proximal straight tubule (PST) in zebrafish embryos at 3 dpf, and in the posterior neural tube at 2 dpf as previously described^52^. Briefly, the embryos were treated with 30 μg/mL 1-phenyl 2-thiourea (PTU) to inhibit pigmentation from 24 hpf. Ciliary motility was measured right after the heartbeat was stopped by incubation with 2,3-butanedione monoxime (BDM). Ciliary movement was recorded using 63× oil objective on an Olympus IX71 microscope equipped with a high-speed camera at a rate of 200 frames per second, and playback set at 20 frames per second. Images were processed using ImageJ software.

### Isolation of renal tubules from zebrafish

For RNA-seq and ATAC-seq, the pronephros were isolated from wild-type or *actl6a^−/−^* embryos with transgenic *Tg (Na,K-ATPase alpha1A4:GFP)* background as previously described^39^. The embryos were dissected, and each pronephroi was collected by visual identification according to GFP fluorescence. The fragments of pronephros were aspirated with an injection needle and transferred into a tube containing lysis buffer for library construction.

### RNA-seq and data analysis

RNA-seq libraries were constructed as previously described^53^. Briefly, kidney tubules from control sibling and *actl6a*-depleted embryos at 30 hpf and 72 hpf were isolated, and total RNA extracted. Sequencing libraries were prepared using the Hyper Prep Kits (KK8504; KAPA Biosystems, USA) according to the manufacturer’s instructions and sequenced on an Illumina Novaseq platform, yielding 20 to 70 million 300-bp reads per sample.

Raw reads from RNA-seq were trimmed with Cutadapt^54^ and quality-controlled with FastQC to remove poly A, TSO, low quality and adaptor-contaminated reads^55^. Clean reads were aligned to the GRCz11 zebrafish genome reference using STAR and quantified with featureCounts^56,57^. Differential expressed genes were identified using edgeR with a threshold of >1.5-fold change and p-value < 0.05^58^. Motif enrichment analysis for ciliary genes was conducted using the iRegulon tool on the down-regulated gene list from 30 hpf and 72 hpf RNA-seq results^59^.

### ATAC-seq experiment

The Hyperactive ATAC-Seq Library Prep Kit for Illumina (Vazyme, China) was utilized to construct ATAC-seq libraries following the manufacturer’s instructions. Briefly, renal tubules from 30 hpf control and *actl6a* morphant embryos were dissociated into single cells. Libraries were prepared from 10,000-20,000 viable cells per sample and sequenced on an Illumina Novaseq platform.

### FitCUT&RUN library preparation

FitCUT&RUN libraries were prepared as previously described^60^. In short, 30 hpf embryos from control and *actl6a* morphants were co-injected with Smarca4a-rFc mRNA and dissociated into single cells (18 µL 0.9 U/µL papain, 2 µL 10x triple LE, 4 µL 1% DNase, 8 µL L-cysteine, 68 µL DMEM/F12, 100 µL 0.25% trypsin). For each sample, 200,000 viable cells were resuspended for library preparation. For FitCUT&RUN, the antibody incubation step was omitted, and libraries were amplified from purified DNA using a Hyper Prep kit (KK8504; KAPA Biosystems, USA). Libraries were sequenced on an Illumina Novaseq.

### ATAC-seq and FitCUT&RUN data analysis

All sequence reads were aligned to the GRCz11 reference genome using Bowtie2 with the parameters “-N 0 -X 2000 --no-mixed --no-unal --no-discordant” after quality trimming with trim-galore (parameters: “--length 20 --paired -q 20 --trim-n”)^61,62^. Data quality was assessed using FastQC^55^. Signal tracks for each sample were generated using the deepTools bamCoverage function with normalization to RPKM (Reads Per Kilobase Million)^63^. Heatmaps of mean read density across all peaks were generated using deepTools. The results (bigWig files) were visualized as track views generated with the Integrative Genomics Viewer (IGV)^64^. To evaluate the reproducibility of the FitCUT&RUN and ATAC-seq experiments, Pearson correlation coefficients were calculated, and principal component analysis was performed on the signals across biological replicates in the merged peaks.

#### Identification of ATAC and FitCUT&RUN peaks

The peaks of control and *actl6a* morphants were identified using MACS2^65^ with the parameter “-g 1.35e9 – nomodel -q 1e-5”. To assess the difference between *actl6a* morphants and control experiments, specific peaks (upregulated and downregulated) were defined using the BedTools subtract function with the parameter “-A”.

#### Annotation and definition

Gene annotation of ATAC and FitCUT&RUN peaks was performed using the Homer annotatePeaks command^66^. The associations between cis-regulatory elements and genes were defined based on the closest peaks and distances to the transcription start sites (TSS) of less than 1 million base pairs.

### Gene ontology analysis

For RNA-seq data, the Database for Annotation Visualization and Integrated Discovery (DAVID; https://david.ncifcrf.gov/)^67,68^ was used to identify Gene Ontology (GO) terms for the biological functions associated with the upregulated and downregulated genes. Genes were tested for enrichment in Gene Ontology Resource (GO) Biological Process (BP), Cellular Component (CC) and Molecular Function (MF) databases.

For ATAC-seq and FitCUT&RUN, functional enrichment analysis of gene sets was performed using the ClusterProfiler tool in R^69^. The 12 functions with the minimum q-values in each gene cluster were selected for comparison against all samples.

Additionally, Metascape software (Metascape; https://metascape.org/)^70^ was utilized to identify GO terms for the biological functions associated with the intersection of different sequencing results.

### Data availability

The raw sequence data from this study have been deposited in the Gene Expression Omnibus (GEO) with accession numbers GSE277562, GSE277564 (RNA-seq), GSE274957 (ATAC-seq), and GSE274953 (FitCUT&RUN). Source data are provided in this paper.

### Statistics

Standard deviations and statistical significance were calculated using GraphPad Prism (GraphPad, USA). The student’s t test was used for analysis. P-values were considered statistically significant at thresholds of P < 0.05, P < 0.01, or P < 0.001.

## Acknowledgments

We thank the members of the Cao lab for helpful discussions. This project was supported by grants from National Key Research and Development Program of China (2017YFA0104600), National Natural Science Foundation of China (32170835 and 31970767).

## Author Contributions

Y. Cao and X. Cheng designed research; X. Cheng, S. Ma, G. Huang, G. Liu and W. Zhang performed research; X. Cheng and Y. Cao analyzed the experimental data; Q. Zhu and X. Peng analyzed the omics data; Y. Cao and X. Cheng wrote the paper; C. Jiang, Y. Zhang and A. Qiu contributed new analytic tools or experiments.

## Declaration of interests

The authors declare no competing interests

---

**Figure S1:**
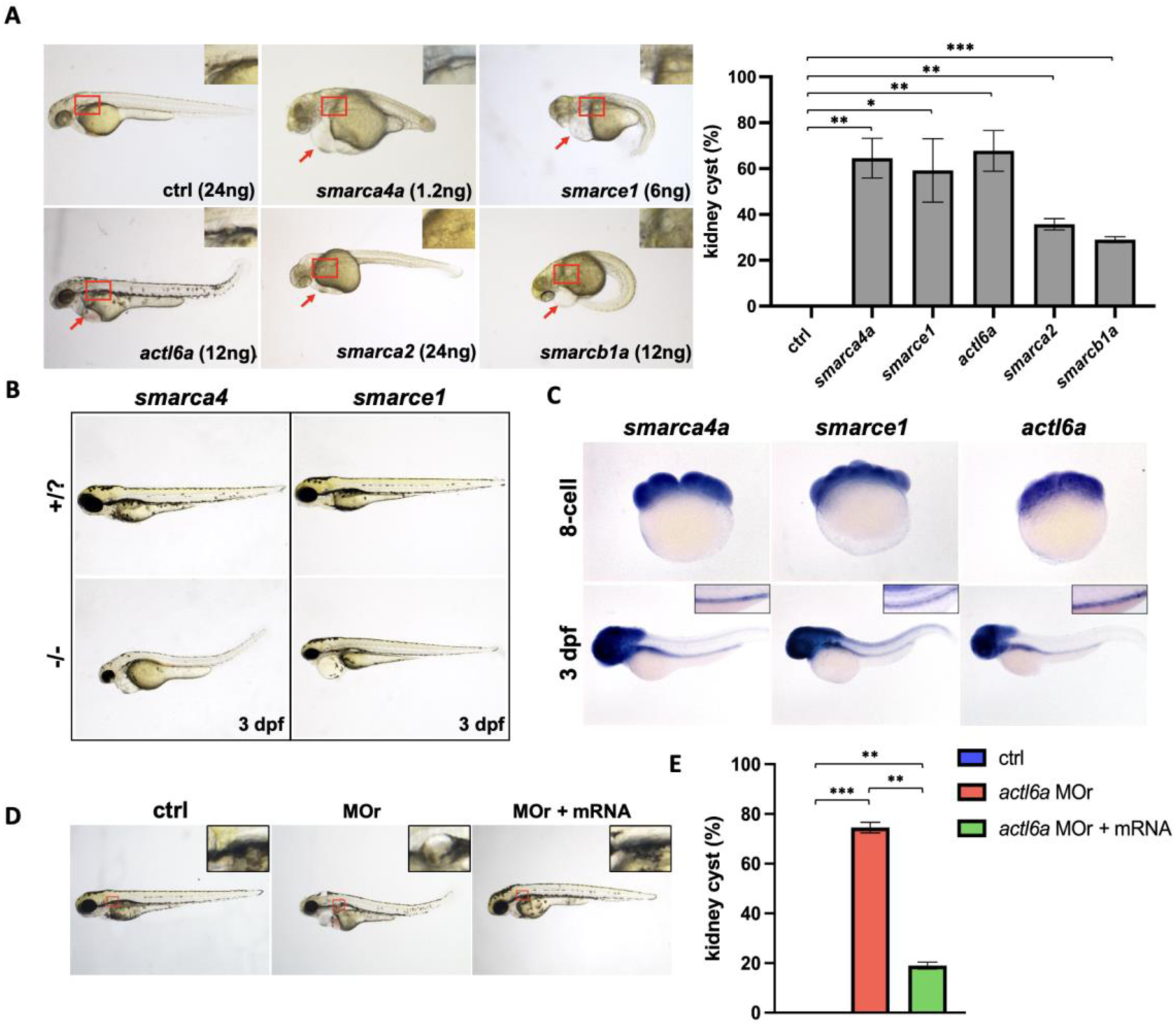
the SWI/SNF chromatin remodeling complex is required for normal kidney development. (A) Representative images of 3 dpf embryos, in which *smarca4a*, *smarce1*, *actl6a*, *smarca2*, or *smarcb1a* were knockdown by injecting respective MOs at 1-cell stage. Insets show a magnified view of the glomerulus area within the boxed regions. Red arrows indicate heart edema. Numbers in parentheses denote the injection doses. Percentage of embryos with kidney cysts were calculated and plotted. n = 50-150 embryos per group, N = 2 repeats. (B) Mutants of *smarca4a* and *smarce1* exhibit heart edema and small eyes at 3 dpf. (C) Whole-mount RNA in situ hybridization of *smarca4a*, *smarce1*, and *actl6a* in wild-type embryos at the 8-cell stage and at 3 dpf. Insets show magnified views of the pronephric tubules in the main panels. (D) Overexpression of *actl6a* mRNA rescues kidney cysts in *actl6a* morphants. Insets show a magnified view of the glomerulus area within the boxed regions. (E) Quantification of the percentage of embryos with cystic kidney in (D). n = 50-150 embryos per group, N = 2 repeats. The data are presented as the mean±SD. \**P* < 0.05, \*\**P* < 0.01, \*\*\**P* < 0.001. Abbreviations: dpf, days post-fertilization; MOs, morpholinos.

**Figure S2:**
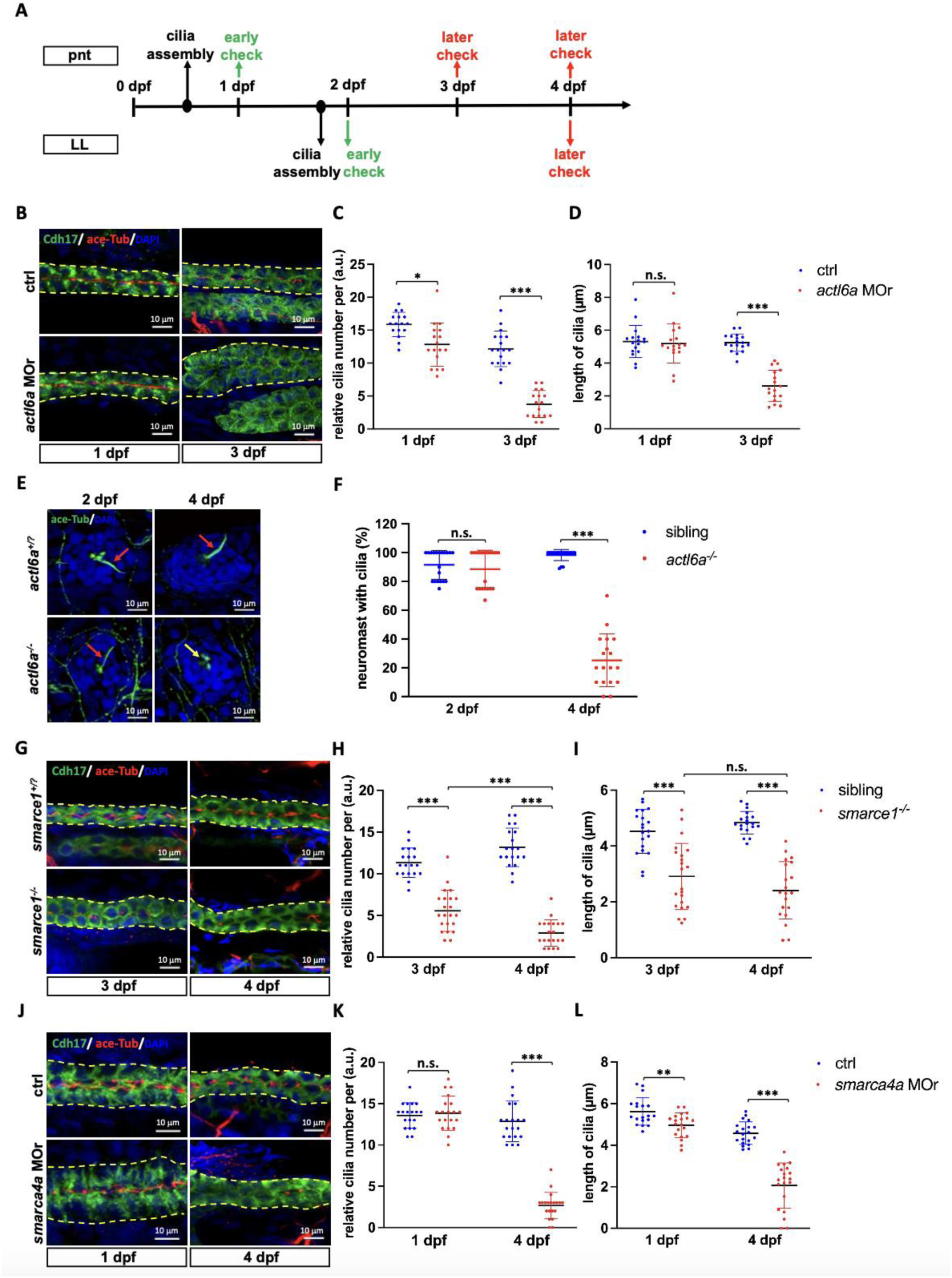
SWI/SNF complex maintains ciliary stability. (A) Schematic illustration of the timepoints for ciliary assembly and detection in the pnt (above the line) or LL (under the line). (B) *actl6a* morphants display normal or disassembled cilia (Acetylated-⍺-Tubulin, red) in the distal pnt (Cdh17, green) at 1 dpf or 3 dpf, respectively. Yellow dotted lines outline the pnt. The number of single cilia per a.u. and cilia length are quantified and plotted in (C-D). n = 8-10 embryos per group, N = 2 repeats. (E) Immunofluorescent images show normal or dissembled cilia (Acetylated-⍺-Tubulin, green) in the neuromast cells of lateral line in *actl6a^−/−^* at 2 dpf or 4 dpf, respectively. Red arrows indicate normal cilia, and the yellow arrows indicate dissembled cilia. The percentage of neuromast cells with cilia are quantified and plotted in (F). n = 8-10 embryos per group, N = 2 repeats. (G) *smarce1^−/−^* embryos display progressive disassembly of single-cilia (Acetylated-⍺-Tubulin, red) in the distal pnt (Cdh17, green) at 3 dpf and 4 dpf. Yellow dotted lines outline the pnt. The number of single cilia per a.u. and cilia length are quantified and plotted in (H-I). n = 8-12 embryos per group, N = 2 repeats. (J) The *smarca4a* morphants shows normal (1 dpf) or disassembled (4 dpf) cilia (Acetylated-⍺-Tubulin, red) in the distal pnt (Cdh17, green) at 1 dpf or 4 dpf, respectively. Yellow dotted lines outline the pnt. The number of single cilia per a.u. and cilia length are quantified and plotted (K-L). n = 8-12 embryos per group, N = 2 repeats. The data are presented as the mean±SD. n.s. not significant, \**P* < 0.05, \*\**P* < 0.01, \*\*\**P* < 0.001. Scale bars, 10 µm. Abbreviations: MOs, morpholinos; pnt, pronephric tubule; dpf, days post-fertilization; LL, lateral line.

**Figure S3:**
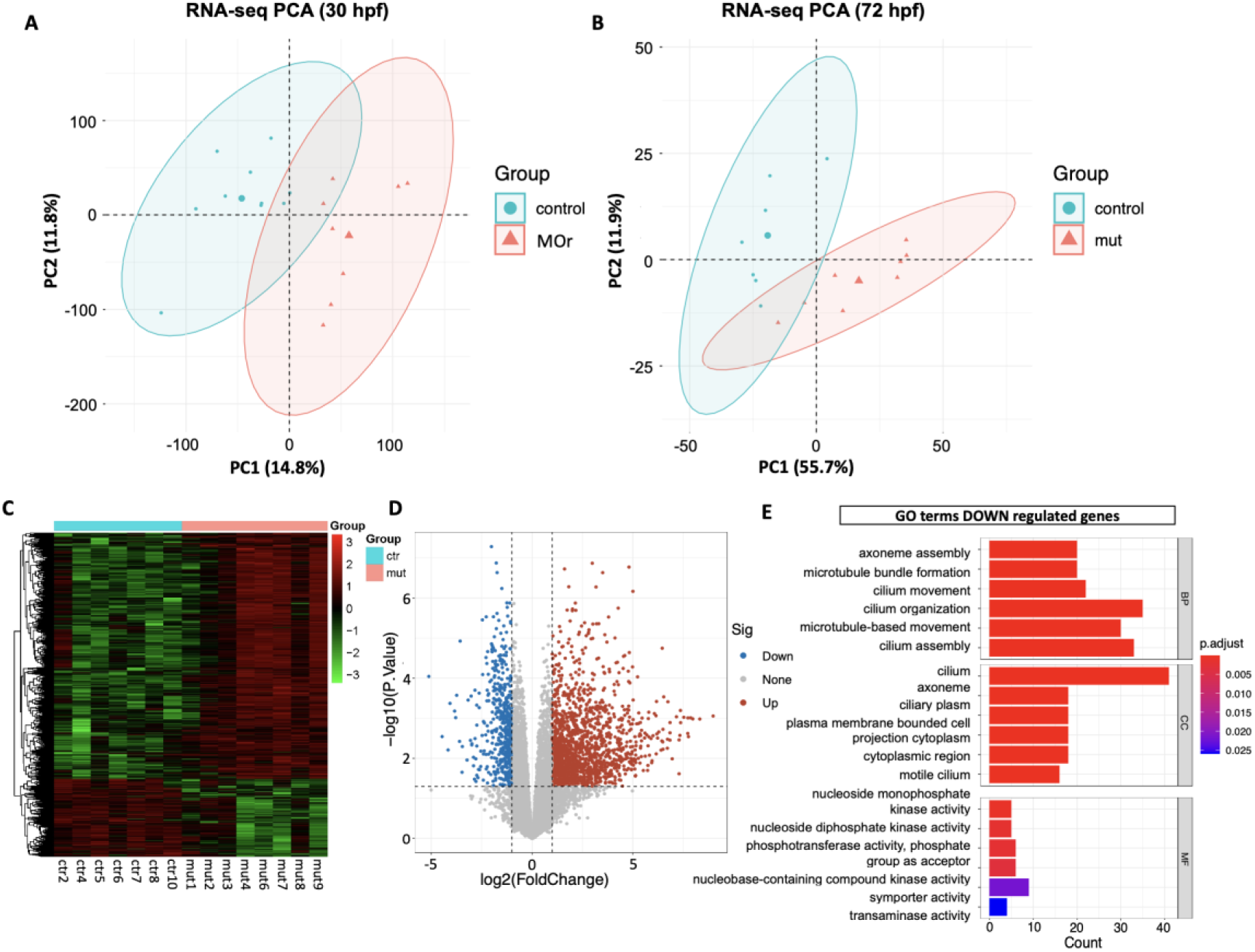
cilia genes are downregulated in the pronephros of *actl6a-*depleted embryos. (A, B) PCA of control and *actl6a*-depleted embryos at 30 hpf (A) or 3 dpf (B) with 95% confidence intervals. (C, D) Heatmap (C) and Volcano plot (D) show differentially expressed genes between control siblings and *actl6a* mutants at 3 dpf. (E) GO analysis of significantly downregulated genes in the *actl6a* mutants compared with the control siblings at 3 dpf. Abbreviations: PCA, principal component analysis; hpf, hours post-fertilization; dpf, days post-fertilization; GO, Gene Ontology.

**Figure S4:**
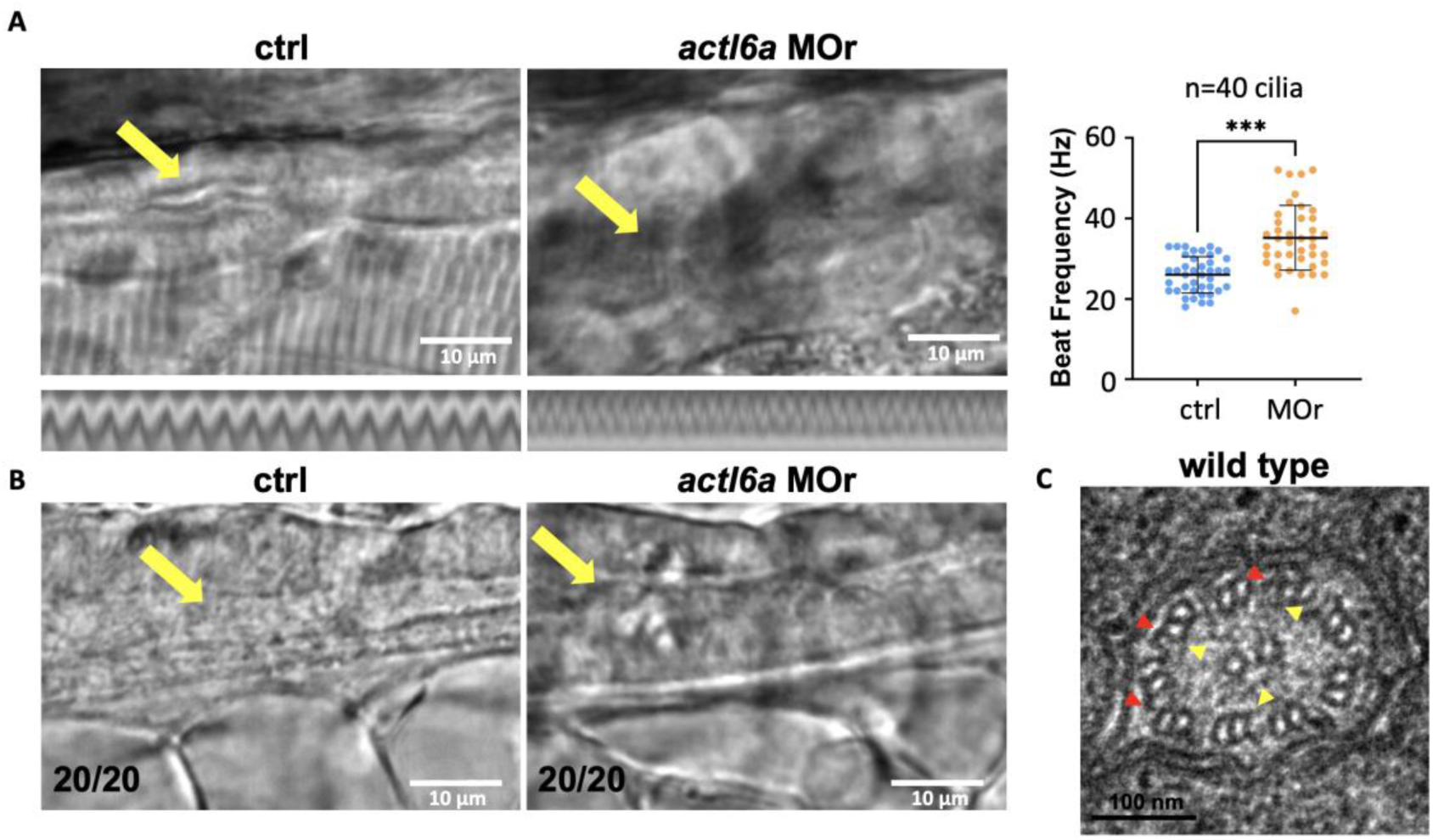
Cilia motility in the pronephric tubule and neural tube remain normal in *actl6a* morphants. (A) Still images show the beating multi-cilia in the middle part of pnt (yellow arrows) in wild-type and *actl6a* morphants at 3 dpf. Representative kymographs of ciliary movement are shown below. The right panel shows statistical analysis of ciliary beating frequency. n = 40 (40 cilia from 8 embryos). Central axis represents the average value. (B) Still images showing the beating cilia in the neural tube (yellow arrows) in wild-type and *actl6a* morphant embryos. n = 10 embryos per group from 2 repeats. (C) Transmission electron micrographs show the “9+2” structure, as well as outer dynein arms (red arrowheads) and inner dynein arms (yellow arrowheads) of single cilia in the distal pnt of wild-type embryos at 3 dpf. The data are presented as the mean±SD. (***) *P* < 0.001. Scale bars, 10 µm in (A, B), 100 nm in (C). Abbreviations: pnt, pronephric tubule; dpf, days post-fertilization.

**Figure S5.**
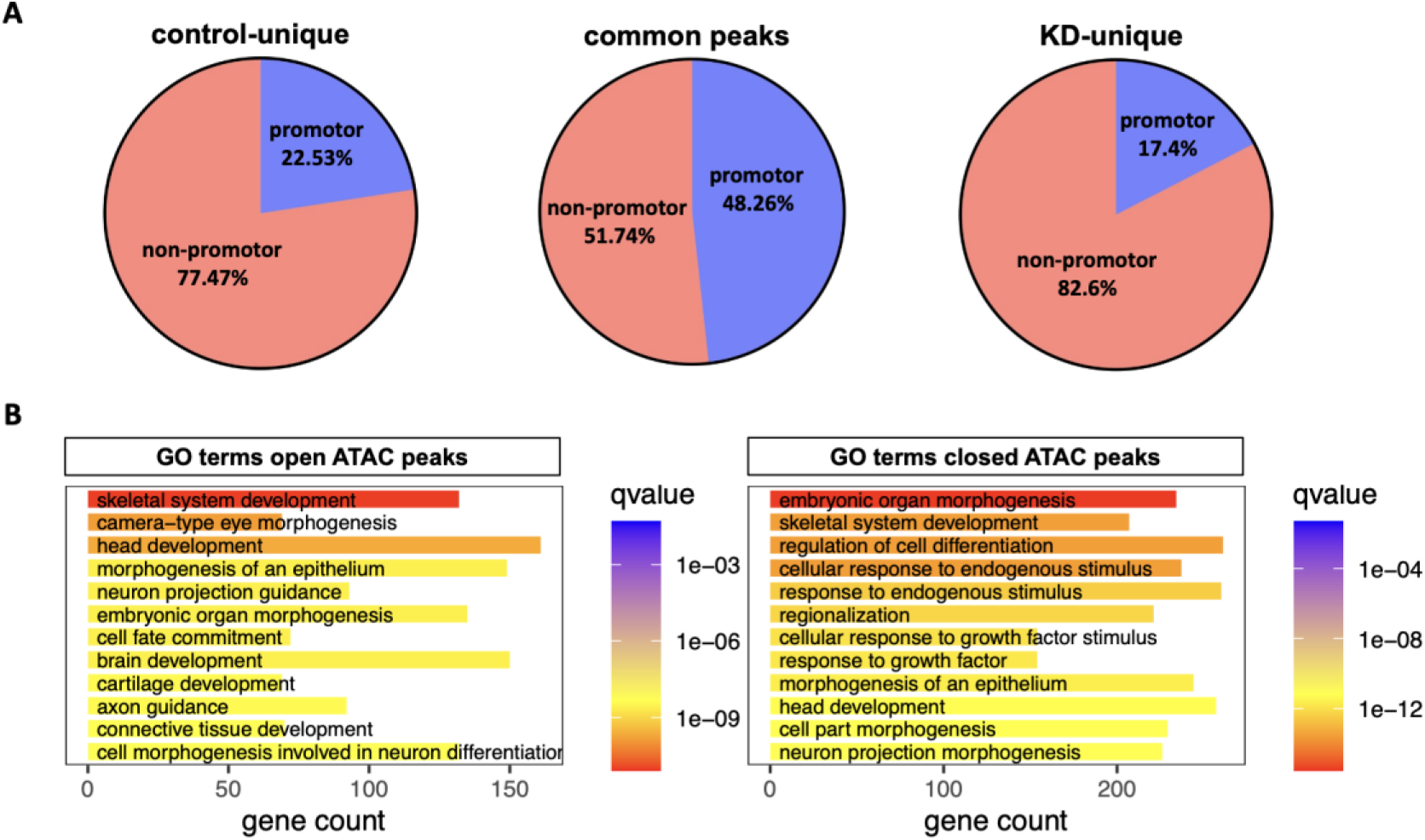
Genomic distribution and GO analysis of ATAC-seq peaks in control and *actl6a*-deficient pronephros. (A) Genomic distribution of accessible ATAC-seq peaks in the control group, with 22.53% located in promoter regions and 77.47% in non-promoter regions. In the *actl6a* morphant group, 17.4% of the peaks are in promoter regions and 82.6% in non-promoter regions. The shared peaks between control and knockdown groups are composed of 48.26% promoter elements and 51.74% non-promoter regions. (B) GO analysis of genes with open (left) and closed (right) chromatin regions in the *actl6a* morphant group compared to the control group, with the top 12 terms shown. Abbreviations: GO, Gene Ontology.

**Figure S6:**
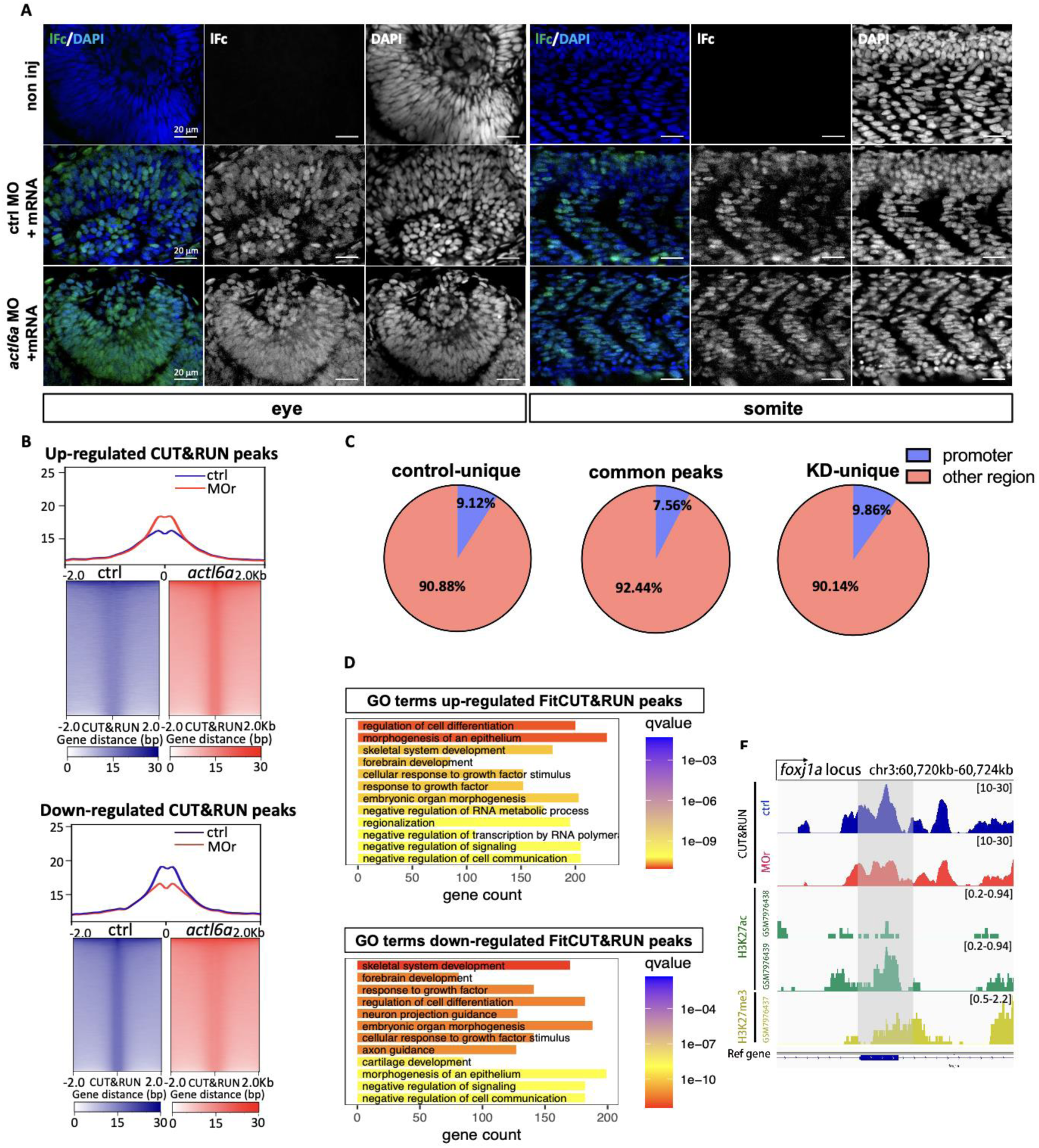
analysis of genomic regions bound by SWI/SNF complex. (A) Immunofluorescent results confirm the validity of rabbit Fc tagged Brg1 (Brg1-rFc) expression. (B) Genome-wide accessibility profiles of FitCUT&RUN at upregulated and downregulated peaks in *actl6a* morphants. (C) Genomic distribution of SWI/SNF complexes binding regions (by FitCUT&RUN) specifically in control or *actl6a* morphants, or common in control and *actl6a* morphants. (D) GO analysis of genes with upregulated or downregulated binding by SWI/SNF complexes in the *actl6a* morphants, with the top 12 terms shown. (E) IGV tracks show downregulated SWI/SNF complexes binding to genomic regions of *foxj1a* in the *actl6a* morphants (red) compared with the control group (blue). The downregulated peak overlaps with enhancer region of *foxj1a* marked by H3K27ac (green) and H3K27me3 (yellow) peaks from a previously published zebrafish dataset. Scale bars, 20 µm. Abbreviations: GO, Gene Ontology; IGV, integrative genomics viewer.

**Figure S7:**
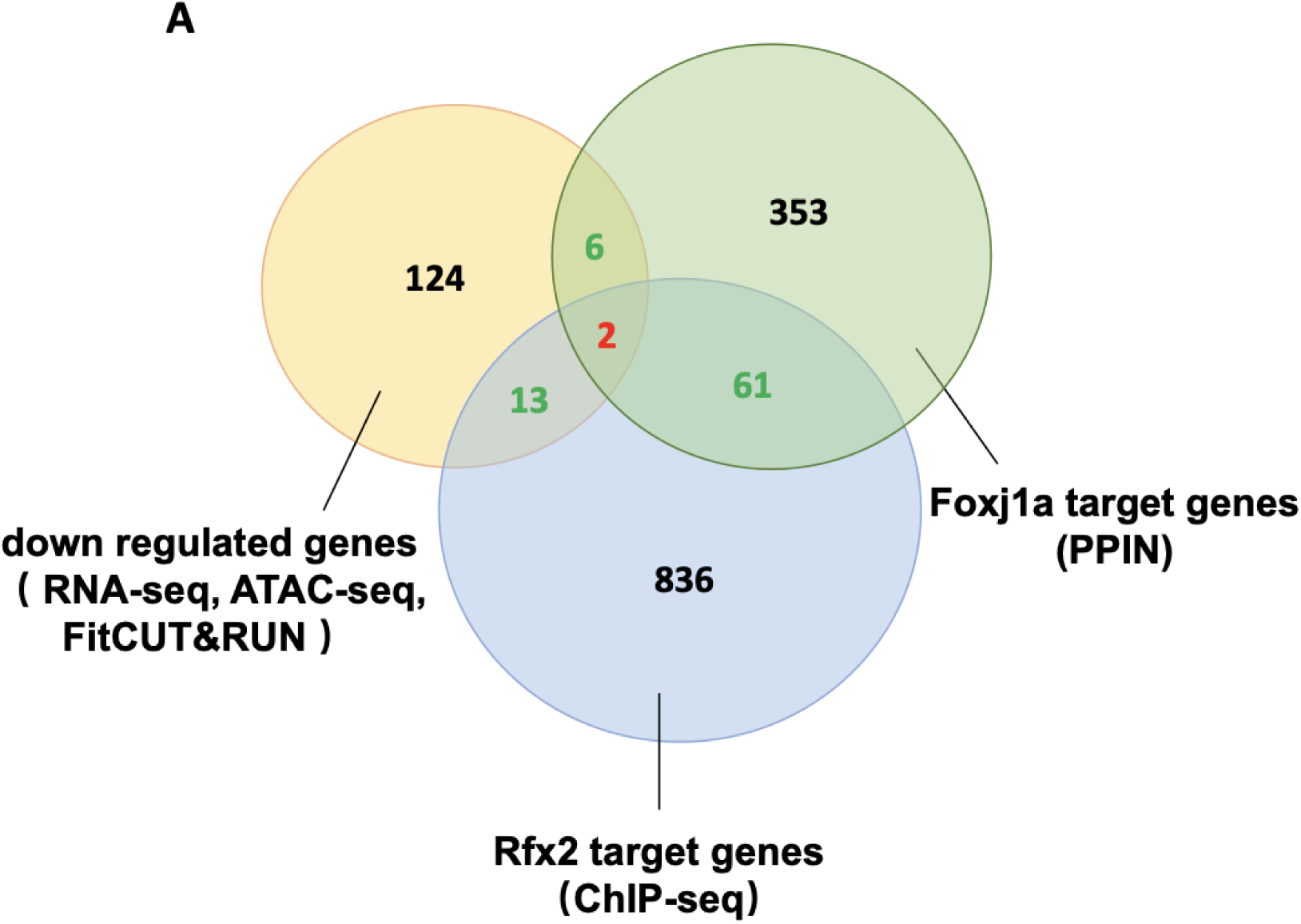
Venn diagram of cilia-related genes directly regulated by *actl6a*, *foxj1a*, and *rfx2*. (A) Venn diagram shows intersection among downregulated genes from three sets of omics data of our study (yellow), Foxj1a direct target genes (green), and Rfx2 direct target genes (blue). The latter two are from previously published data.

**Figure S8:**
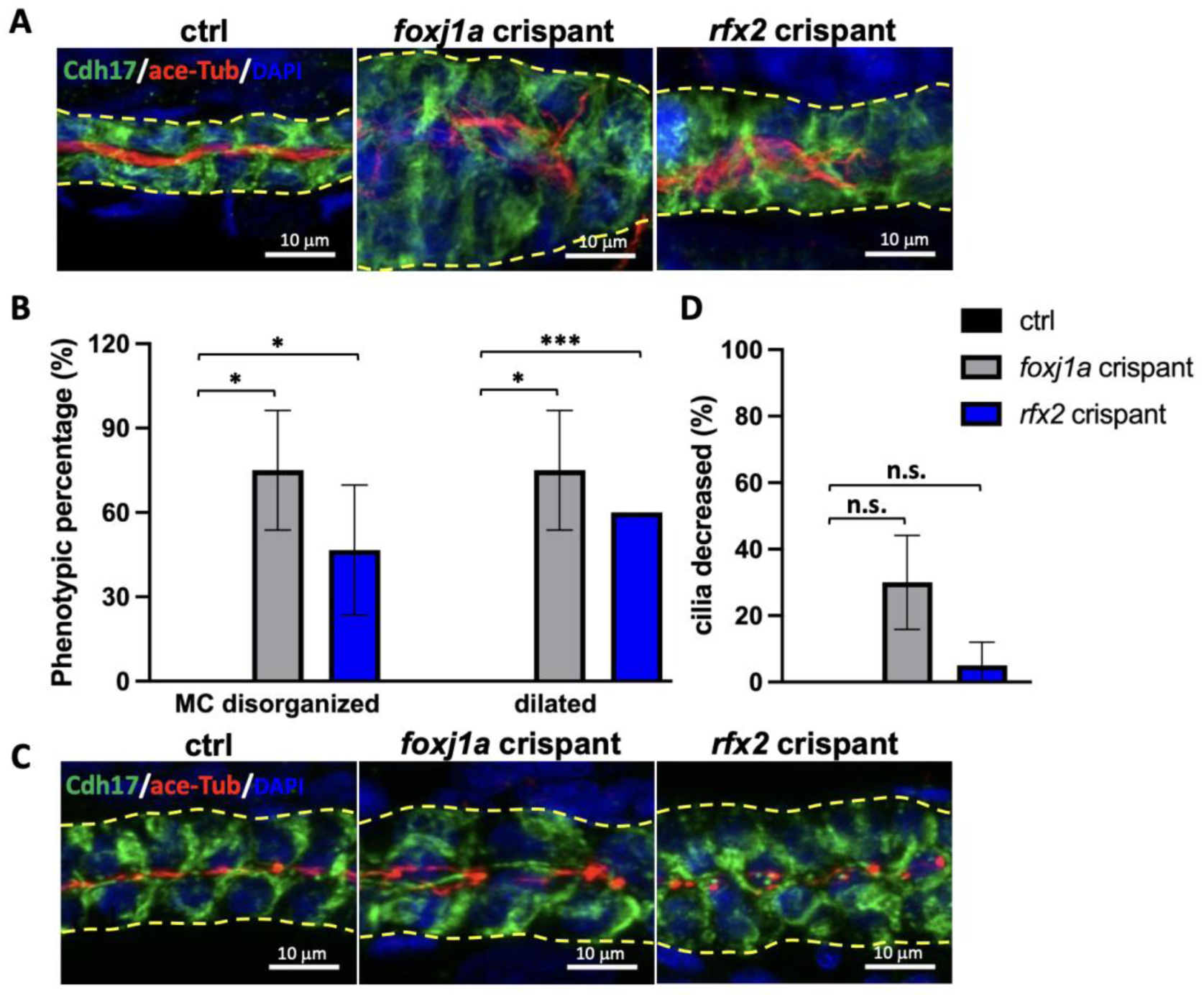
Foxj1a and Rfx2 may not be required for single cilia assembly in the pronephros of zebrafish embryos. (A) At 3 dpf, both *foxj1a* crispants and *rfx2* crispants exhibit dilated pnt (Cdh17, green) and disorganized multi-cilia (Acetylated-⍺-Tubulin, red) in the middle pnt. Yellow dotted lines outline the pnt. (B) The percentage of embryos with dilated pnt and disorganized multi-cilia in *foxj1a* and *rfx2* crispants is quantified and plotted. n = 8-10 embryos per group, N = 2 biological repeats. (C) At 1 dpf, both *foxj1a* crispants and *rfx2* crispants show normal cilia assembly (Acetylated-⍺-Tubulin, red) in the distal pnt (Cdh17, green). Yellow dotted lines outline the pnt. (D) The percentage of embryos with decreased single cilia is quantified and plotted. n = 8-10 embryos per group, N = 2 repeats. The data are presented as the mean±SD. n.s. not significant, \**P* < 0.05, \*\*\**P* < 0.001. Scale bars, 10 µm. Abbreviations: dpf, days post-fertilization; pnt, pronephric tubule.

**Movie 1. Motile cilia in the middle pnt of the control sibling embryos.**

Motile cilia in the middle pnt of control sibling embryos at 3 dpf, captured at 200 frames per second. The portion shown here is 1 second of footage slowed down to 20 frames per second. The cilia beat at an average frequency of 35.3±8.9 Hz. Scale bars, 10 µm.

**Movie 2. Motile cilia in the middle pnt of the *actl6a* mutant embryo.**

Motile cilia in the middle pnt of *actl6a^−/−^* embryos at 3 dpf, captured at 200 frames per second. The images were acquired and processed identically to those in Movie 1. The cilia beat at an average frequency of 35.7±7.3 Hz. Scale bars, 10 µm.

**Movie 3. Motile cilia in the middle pnt of the control morphant embryos.**

Motile cilia in the middle pnt of control embryos (1 mM control MO) at 3 dpf, captured at 200 frames per second. The portion shown here is 1 second of footage slowed down to 20 frames per second. The cilia beat at an average frequency of 26.0±4.5 Hz. Scale bars, 10 µm.

**Movie 4. Motile cilia in the middle pnt of the *actl6a* morphant embryos.**

Motile cilia in the middle pnt in *actl6a* morphants (1 mM *actl6a* MO) at 3 dpf, captured at 200 frames per second. The images were acquired and processed identically to those in Movie 3. The cilia beat at an average frequency of 35.2±8.0 Hz. Scale bars, 10 µm.

**Movie 5. Motile cilia in the posterior part of the neural tube in control morphant embryos.**

Motile cilia in the posterior neural tube of control embryos (1 mM control MO) at 2 dpf, captured at 200 frames per second. The portion shown here is 1 second of footage slowed down to 20 frames per second. n = 20. Scale bars, 10 µm.

**Movie 6. Motile cilia in the posterior part of the neural tube in the *actl6a* morphant embryos.**

Motile cilia in the posterior neural tube of *actl6a* morphants (1 mM *actl6a* MO) at 2 dpf, captured at 200 frames per second. The portion shown here is 1 second of footage slowed down to 20 frames per second. n = 20. Scale bars, 10 µm.

**Supplementary table 1.**

List of DEG and GO analysis of the DEG in *actl6a*-depleted pronephros (RNA-seq) and the enriched motif in down-regulated cilia genes. Fold change >1.5, pvalue < 0.05. DEG, differentially expressed genes.

**Supplementary table 2.**

List of downregulated/closed/unconjugated genes from RNA-seq, FitCUT&RUN and ATAC-seq in *actl6a* morphant group compared to the control group at 30 hpf, and the GO analysis results of this gene set.

**Supplementary table 3.**

List of gRNA and primers for genotype identification.

